# Sec61 translocon inhibitor Flavitransin blocks selectively dengue virus polyprotein insertion into the ER membrane with pan-flavivirus antiviral potency

**DOI:** 10.1101/2025.03.29.646078

**Authors:** Marijke Verhaegen, Marianne Croonenborghs, Nidhi Sorout, Andrea Sartori, Becky Provinciael, Eef Meyen, Joren Stroobants, Enno Hartmann, Volkhard Helms, Kurt Vermeire

**Affiliations:** Laboratory of Molecular, Structural and Translational Virology, Rega Institute for Medical Research, Department of Microbiology, Immunology and Transplantation, KU Leuven, B-3000 Leuven, Belgium; Center for Bioinformatics, Saarland University, Saarbrücken, Saarland, Germany; Department of Food and Drug, University of Parma, 43124 Parma, Italy; Centre for Structural and Cell Biology in Medicine, Institute of Biology, University of Lübeck, 23562 Lübeck, Germany

**Keywords:** Dengue virus, Zika virus, Sec61 translocon, co-translational translocation, antiviral, cyclotriazadisulfonamide

## Abstract

Flavivirus infections by Dengue and Zika virus impose a significant healthcare threat worldwide. At present no FDA-approved specific antiviral treatment is available, and the safety of a vaccine against Dengue virus is still under debate. Here, we report the identification of the CADA derivative flavitransin (FT), with potent activity against DENV serotype 2 in various cell types. Moreover, FT showed consistent anti-flaviviral activity against all four DENV serotypes, and also against Zika and Yellow fever virus. Viral polyprotein biogenesis was completely abolished by FT treatment of DENV-infected cells. Drug profiling by a-time-of-drug-addition assay revealed a post-entry antiviral effect of FT, in line with its anticipated Sec61 inhibitory effect. Subsequent analysis of the individual viral proteins in transfected HEK293T cells indicated that FT suppresses the expression of the structural proteins (pre-membrane and envelope) only. Furthermore, cell free *in vitro* protein translation analysis demonstrated a direct inhibitory effect of FT on the co-translational translocation of the DENV polyprotein across the ER membrane. More specifically, FT inhibited the initiation of protein translocation into the ER that relies on the N-terminal transmembrane region of the capsid subunit of the DENV polyprotein, resulting in rerouting of the viral pre-protein to the cytosol for proteasomal degradation. Finally, selection and genotyping of FT-resistant HCT116 cells revealed a unique A70V mutation in the Sec61α subunit that conferred resistance to FT in infected cells. Long-term exposure of DENV to FT demonstrated a high barrier to resistance development. In conclusion, our data demonstrate that FT selectively interferes with the initiation of ER co-translational translocation of the DENV polyprotein and confirm the critical role of this translocation process in the flavivirus replication cycle.

## INTRODUCTION

Dengue virus (DENV), the causative agent of dengue disease, is a quickly spreading mosquito-transmitted human pathogen, posing a threat to the world’s population. Symptomatic infections can manifest from self-limited DENV fever to a hemorrhagic fever, and eventually to a fatal plasma leakage-induced shock syndrome. In 2023, more than 6.5 million dengue cases and 7,300 dengue-related deaths were recorded. Although the four DENV serotypes primarily co-circulate in (sub)tropical regions, an increasing number of dengue cases have been reported in Europe and North America due to the spread of the *Aedes* mosquito vector in these (thus far) non-endemic areas [1, 2].

Despite the significant health burden, no specific antiviral treatment for DENV exists, and approved vaccines provide only partial protection, underscoring the urgent need for new antiviral strategies. Several drugs with significant antiviral activity against DENV in preclinical studies, including balapiravir, chloroquine, lovastatin and celgosivir, have failed to show efficacy in lowering viral load or achieving favorable clinical outcomes in clinical trials [3]. The small molecule JNJ-641802, which blocks the formation of the viral NS3-NS4B complex, demonstrated favorable pharmacokinetics and safety profile in phase I clinical trials, but its phase II study was recently discontinued [4–6]. Also, the endoplasmic reticulum membrane (ER) α-glucosidase inhibitor hydrochloride UV-4B was reported safe in phase 1 study, but no follow-up clinical studies have been initiated so far [7].

DENV belongs to the genus *of Flavivirus* (family *Flaviviridae*) which share a positive single-stranded RNA genome (**Fig. 1A**). Following its release into the cytoplasm, the viral RNA genome is translated into a single polytopic protein comprising the three structural proteins [capsid (Ca), membrane (M) and envelope (E) protein] of the viral particle and seven non-structural proteins (NS) which are involved in RNA transcription and virus replication [8]. Genome-wide screens identified the ER as a key organelle throughout the viral life cycle which supports viral protein translation, processing, maturation and replication of the viral RNA [9–15]. The flavivirus polyprotein contains 18 transmembrane domains (TMDs) that, similar to cellular (multi-pass) transmembrane proteins, require insertion into the ER membrane (**Fig. 1A**). Therefore, the nascent viral polyprotein is likely targeted towards and incorporated into the ER membrane by the cellular process of co-translational translocation across the Sec61 translocon channel and associated ER membrane protein complex (EMC) [8, 16–18]. For flaviviruses, this process is believed to be directed by a signal peptide (SP)-like sequence within the N-terminal region of the polyprotein. The TMD of the Ca protein (**Fig. 1A**; purple striped TMD) serves as such an ER localization element, guiding the stalled ribosome/nascent chain complex to the Sec61 translocon. Once docked onto the translocon, the stalled ribosome resumes translation, providing the energy for the viral polytopic protein to be inserted into the ER membrane with the prM, E, NS1 and some extended stretches of NS2A, NS2B, NS4A and NS4B proteins exposed to the ER lumen, and the Ca, NS3 and NS5 proteins positioned at the cytoplasmic side of the ER membrane (**Fig. 1A**). The polyprotein is further processed into the separate viral proteins by a cellular signal peptidase in the ER lumen and by the viral protease (i.e., NS3 and its cofactor NS2B) in the cytosol. The C-terminal TMD of the Ca protein, the prM and the E protein, and the 2K peptide separating NS4A and NS4B, all contain a proteolytic cleavage site that is recognized by the signal peptidase, hence, these TMDs are designated as SP-like targeting sequences for the downstream viral subunits (prM, E, NS1 and NS4B, respectively) (**Fig. 1A**) [11, 19–24]. ER-resident enzyme complexes (such as oligosaccharyltransferase complex) and chaperones regulate the N-glycosylation and folding of the viral proteins, facilitating their maturation [25–29]. The NS proteins reside at the ER to orchestrate the replication complexes for the synthesis of new viral RNA copies. As a final step at the ER, the viral structural proteins assemble and bud with incorporation of the newly synthesized RNA to form individual virus particles [30]. These immature virions are transported via the cellular secretory pathway, undergoing final maturation (e.g., pH-dependent furin cleavage) at the Golgi complex [31].

**Figure 1.**
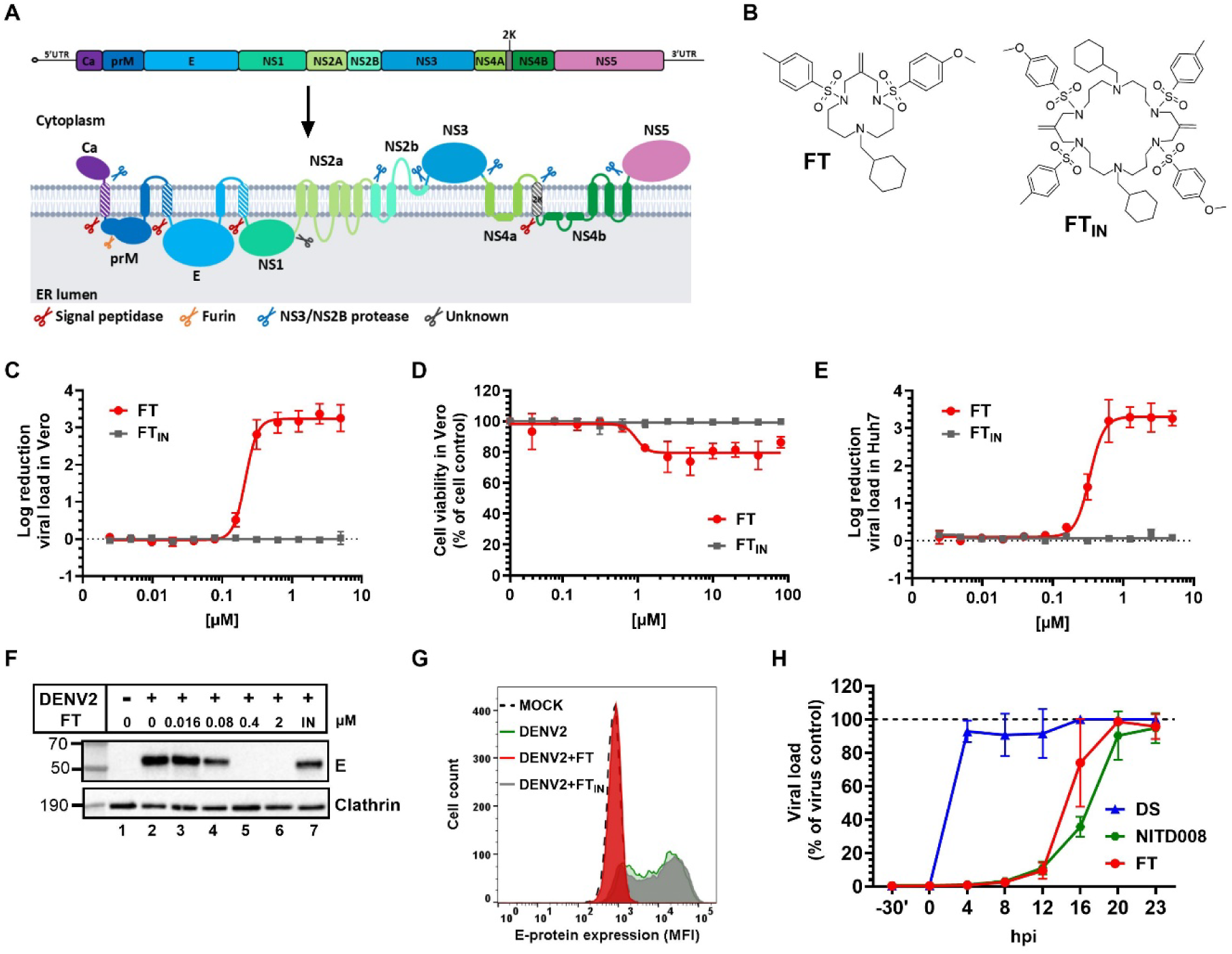
Flavitransin inhibits DENV2 replication at a post-entry event. **A**, Schematic representation of a flaviviral polyprotein embedded in the ER membrane with 18 membrane-spanning domains. The different proteolytic cleavage sites are indicated by colored scissors. The viral protease, formed by NS3 and its cofactor NS2B (blue scissors), releases the viral subunits at the cytoplasmic side of the ER membrane. Shaded transmembrane domains (boxes) indicate cleavable SP-like targeting sequences for Sec61 translocation. **B**, Chemical structure of FT and FT_IN_. **C**, Four parameter concentration-response curve of FT or FT_IN_ for virus yield (log_10_ reduction in copy number compared to untreated virus control) in DENV2 NGC-infected (MOI 1) Vero cells. Compound treatment started 2 h before infection, and supernatant was collected at 4-5 days post infection for RT-qPCR detection of the viral 3’UTR. Data points show mean values ± SD, n=3. **D**, Four parameter concentration-response curve of FT or FT_IN_ for viability of Vero cells. Cytotoxic effects were determined at day 5 post treatment by MTS/PMS assay. Data are normalized to untreated cell controls. Graph shows mean values ± SD, n=3. **E**, Same as in (C) but for Huh7 cells. **F**, Viral E protein detection in DENV2 NGC (MOI 0.3) infected Huh7 cells, treated with FT or 10 µM FT_IN_ (indicated as ‘IN’) for 48 h. Immunoblot on cell lysate with anti-E or anti-clathrin (loading control) antibody. **G**, Viral E protein detection in DENV2 NGC (MOI 0.5) infected Huh7 cells, treated with 2 µM FT or 10 µM FT_IN_ for 72h. Flow cytometric histogram plots show mean fluorescence intensity (MFI) values for E protein, acquired from at least 5,000 cells. Mock-infected cells are in black dotted line, virus infected control cells are in green line, FT treated infected cells in red and FT_IN_ infected cells in grey. Representative histogram plots are given for one out of two experiments. **H**, Time-of-drug-addition assay in which Vero cells were treated with the attachment inhibitor Dextran Sulfate (DS, 4 mg/ml), FT (1 µM) or the viral polymerase inhibitor NITD008 (10 µM) starting before infection (-30’), at the time of infection with DENV2 NGC (0 h) or at several time points after infection (4, 8, 12, 16, 20, 23 h) . Cells were exposed to virus for 2 h and excess of virus was then removed. Compounds were present till end of experiment. At 24 hpi, cells were lysed for viral RNA quantification by RT-qPCR (3’UTR). Graphs represent the viral load as normalized to the viral copy number of the virus control from 3 independent experiments. Values are mean ± SD; n=3.

Numerous components of the cellular Sec61 co-translational translocation pathway have been identified as important host factors for flavivirus replication [9–13]. The heterotrimeric Sec61 complex is formed by Sec61α, Sec61β, and Sec61γ subunits that span the eukaryotic ER membrane [32]. The Sec61α subunit consists of ten transmembrane helices (TMHs) that surround the central pore of the translocon and their structural rearrangements regulate the different steps of protein translocation. The central pore is closed by a movable plug domain at the ER luminal side and sealed by the lateral gate, formed by TMH2, 3, 7 and 8, in its native state. Binding of the ribosome nascent chain complex (primed state) and interaction of the targeting sequence (e.g., SP) with the translocon interrupts the interhelical contacts between the TMH2 and TMH7 of the lateral gate (SP engaged state). As translation progresses, the inserting polypeptides either laterally insert into the ER membrane by further opening of the lateral gate (lateral insertion) or translocate along the Sec61 pore into the ER lumen by displacement of the plug domain (vertical translocation) [33, 34]. The hydrophobic strength of the targeting sequence or TMDs of the nascent protein chains is a crucial parameter for successful protein translocation. Only sufficiently hydrophobic targeting peptides succeed to displace the plug domain or disrupt the interhelical hydrophobic interaction between the TMHs to open the lateral gate. Proteins with weak hydrophobic targeting sequences or TMDs require accessory components such as the EMC, translocon-associated protein (TRAP) complex, translocating chain-associated membrane protein (TRAM), Sec62, Sec63 and BiP for ER translocation [35, 36].

Although completely blocking protein import via Sec61 is likely toxic to cells, there is significant interest in identifying small-molecule inhibitors of Sec61 that may either only block protein import selectively, or that may be used at lower, sub-toxic concentrations. Hence, inhibitors of Sec61-dependent protein translocation are considered promising candidates for anti-cancer and immunosuppressive therapies and to treat several human diseases [35, 37, 38]. However, their potential in the context of antiviral treatment has received limited attention. The Sec61 translocon inhibitors cotransin 8 and its analogue PS3061 have been shown to effectively suppress DENV replication in monocyte-derived dendritic cells and vector cells [13, 39]. Also, Apratoxin S4 exerted a profound pan-flaviviral activity against DENV2, Zika virus (ZIKV), and West Nile virus (WNV) [40]. Recently, the highly potent but toxic Sec61 inhibitor mycolactone was shown to block the production of virus glycoproteins of a range of enveloped viruses, including ZIKV [41].

The small synthetic molecule cyclotriazadisulfonamide (CADA) was originally discovered as an anti-human immunodeficiency virus (HIV) agent that downmodulates the cell surface expression of human cluster of differentiation 4 (huCD4), the main entry receptor of HIV that initiates the fusion of the viral envelope with the cell membrane. The antiviral effect of CADA can thus be explained by the reduction in surface levels of huCD4 to levels below the threshold that is required for HIV entry into the host cell [42]. We previously established that CADA suppresses huCD4 cell surface expression via SP-dependent inhibition of the co-translational translocation of the huCD4 pre-protein into the ER [43]. Due to the relatively small ring structure of CADA, its chemical synthesis is feasible and straight forward, allowing for the design and generation of a library of structural analogs to improve the potency and selectivity of the compound. From this library, compound CK147 was selected as the most potent unsymmetrical CADA analog discovered so far with a nearly 10-fold enhanced potency as compared to CADA [44], and was used to resolve a high-resolution single-particle cryo-EM structure of the Sec61 translocon in complex with a non-translating ribosome and a Sec61 inhibitor [45].

In our current study, we identified the CADA derivative Flavitransin (FT; **Fig. 1B**) as a promising hit with pan-DENV serotype and pan-flavivirus antiviral potency in the sub-micromolar range. We further demonstrated that FT selectively inhibits Sec61-directed translocation of the viral polytopic preprotein and diverts the viral nascent chain to the cytosol for proteasomal degradation. *In vitro* translocation assays revealed that FT selectively blocks the insertion of the first translated N-terminal TMD of the DENV polytopic protein into the ER membrane. Finally, FT-resistant cells display unique mutations in the Sec61α translocon subunit.

## RESULTS

### Flavitransin is a potent inhibitor of DENV2 infection

An in-house antiviral screen against Dengue virus (DENV) identified a hit compound with high nanomolar activity. This small molecule, designated as Flavitransin (FT; **Fig. 1B**), inhibited concentration-dependently DENV2 replication in African green monkey kidney cells (Vero), whereas FT_IN_, a double size ring structured analogue (obtained during the chemical synthesis of FT as a minor side-product fraction) had no antiviral effect (**Fig. 1C**; **Table 1**). FT completely blocked DENV2 infection of Vero cells as determined by RT-qPCR (3’UTR of the viral genome; **Fig. 1C**) and E protein measurement (**Suppl. Fig. 1A**), with low impact on cell viability (**Fig. 1D**). Similar antiviral effect of FT was observed in human hepatocarcinoma (Huh7) cells, as evidenced by viral copy number measurement (**Fig. 1E**) and E protein quantification (with immunoblotting and flow cytometry; **Fig. 1F, G** and **Suppl. Fig. 1B; Table 1**). Moreover, 0.4 µM FT treatment of DENV2 infected Huh7 cells completely suppressed the cellular expression of all viral structural and NS proteins, whereas a 10 µM FT_IN_ exposure did not (**Fig. 1F** and **Suppl. Fig. 2**).

**Table 1.**
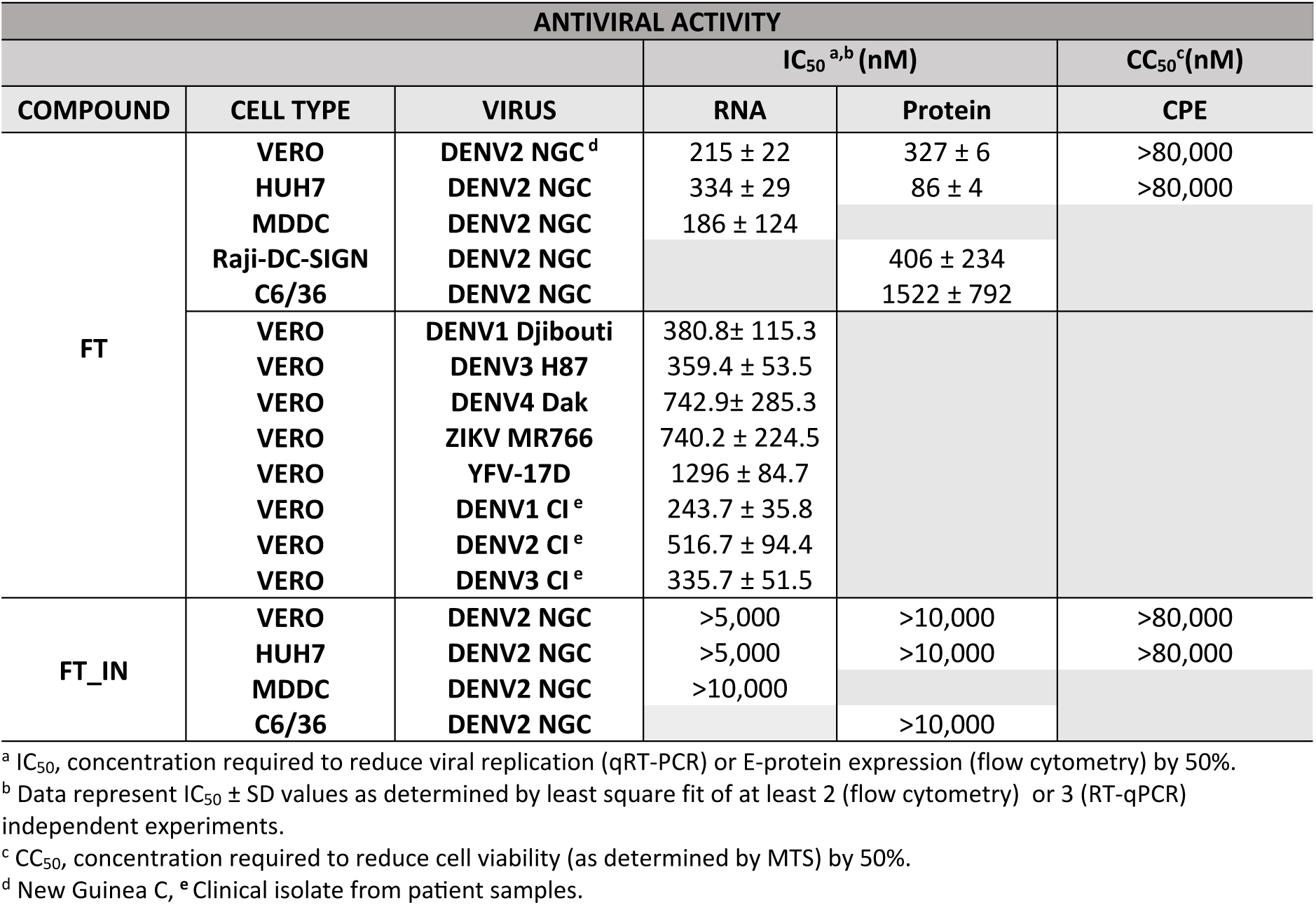
Antiviral activity and toxicity of FT and FT_IN_ against flavivirus in different cell types.

| ANTIVIRAL ACTIVITY |  |  |  |  |  |
| --- | --- | --- | --- | --- | --- |
|  |  |  | IC<sub>50</sub><sup>a,b</sup> (nM) |  | CC<sub>50</sub><sup>c</sup> (nM) |
| COMPOUND | CELL TYPE | VIRUS | RNA | Protein | CPE |
| FT | VERO | DENV2 NGC<sup>d</sup> | 215 ± 22 | 327 ± 6 | >80,000 |
|  | HUH7 | DENV2 NGC | 334 ± 29 | 86 ± 4 | >80,000 |
|  | MDDC | DENV2 NGC | 186 ± 124 |  |  |
|  | Raji-DC-SIGN | DENV2 NGC |  | 406 ± 234 |  |
|  | C6/36 | DENV2 NGC |  | 1522 ± 792 |  |
|  | VERO | DENV1 Djibouti | 380.8 ± 115.3 |  |  |
|  | VERO | DENV3 H87 | 359.4 ± 53.5 |  |  |
|  | VERO | DENV4 Dak | 742.9 ± 285.3 |  |  |
|  | VERO | ZIKV MR766 | 740.2 ± 224.5 |  |  |
|  | VERO | YFV-17D | 1296 ± 84.7 |  |  |
|  | VERO | DENV1 CI<sup>e</sup> | 243.7 ± 35.8 |  |  |
|  | VERO | DENV2 CI<sup>e</sup> | 516.7 ± 94.4 |  |  |
|  | VERO | DENV3 CI<sup>e</sup> | 335.7 ± 51.5 |  |  |
| FT<sub>IN</sub> | VERO | DENV2 NGC | >5,000 | >10,000 | >80,000 |
|  | HUH7 | DENV2 NGC | >5,000 | >10,000 | >80,000 |
|  | MDDC | DENV2 NGC | >10,000 |  |  |
|  | C6/36 | DENV2 NGC |  | >10,000 |  |

<sup>a</sup> IC<sub>50</sub>, concentration required to reduce viral replication (qRT-PCR) or E-protein expression (flow cytometry) by 50%.
<sup>b</sup> Data represent IC<sub>50</sub> ± SD values as determined by least square fit of at least 2 (flow cytometry) or 3 (RT-qPCR) independent experiments.
<sup>c</sup> CC<sub>50</sub>, concentration required to reduce cell viability (as determined by MTS) by 50%.
<sup>d</sup> New Guinea C, <sup>e</sup> Clinical isolate from patient samples.

### Flavitransin interferes with a post entry event of the DENV2 life cycle

To further profile FT, we determined by means of postponing treatment in a time-of-drug-addition assay at what stage during the viral life cycle FT exerts its inhibitory effect. Briefly, the compound was administered to DENV2 infected Vero cells either before or at specific time points after virus exposure, and viral copy numbers in infected cells were determined at 24 h post infection, roughly when one viral life cycle has been completed (**Suppl. Fig. 3A**). We observed a clear post entry effect of FT, given that the administration of the drug could be delayed by approximately 8 h with preservation of its antiviral effect (**Fig. 1H**), whereas the attachment inhibitor Dextran Sulfate (DS) was only effective when administered before or at the time point of virus exposure. The profile of FT resembled that of the replication inhibitor NITD008 (**Fig. 1H**), a nucleoside analogue that inhibits the viral polymerase of DENV [46]. Together, these data suggest that FT interferes with a virus replication event taking place between 4-8 hpi. Given the structural resemblance of FT with the earlier reported translocon inhibitors CADA and CK147 [43, 45], translocation of the viral protein across the ER membrane was considered a plausible target of FT and subject for further investigation.

### Flavitransin inhibits the expression of structural proteins of DENV2 in transfected cells

To examine an inhibitory effect of FT on the translation and/or processing of the DENV polytopic protein, we first dissected the impact of FT on the cellular expression of individual DENV proteins from single expression vectors. We focused on the protein subunits prM-E and NS1 as these are preceded by a cleavable signal peptide-like hydrophobic sequence for ER Sec61 targeting (**Fig. 2A**). The prM/E precursor contains 2 proteolytic cleavage sites (recognized by signal peptidases in the ER lumen; **Fig. 1A**) for further processing into the prM and E structural subunits. Transfection of HEK293T cells with this construct resulted in successful protein expression in the non-treated samples, giving rise to prM and E species of approximately 20 and 55 kDa, respectively (**Fig. 2B**, lane 2), similar to what was detected in DENV2 infected Huh7 cells (**Fig. 1F** and **Suppl. Fig. 2**). Importantly, treatment of the transfected cells with FT resulted in a significant concentration-dependent inhibition of both prM and E expression (**Fig. 2B**, lanes 3-5).

**Figure 2.**
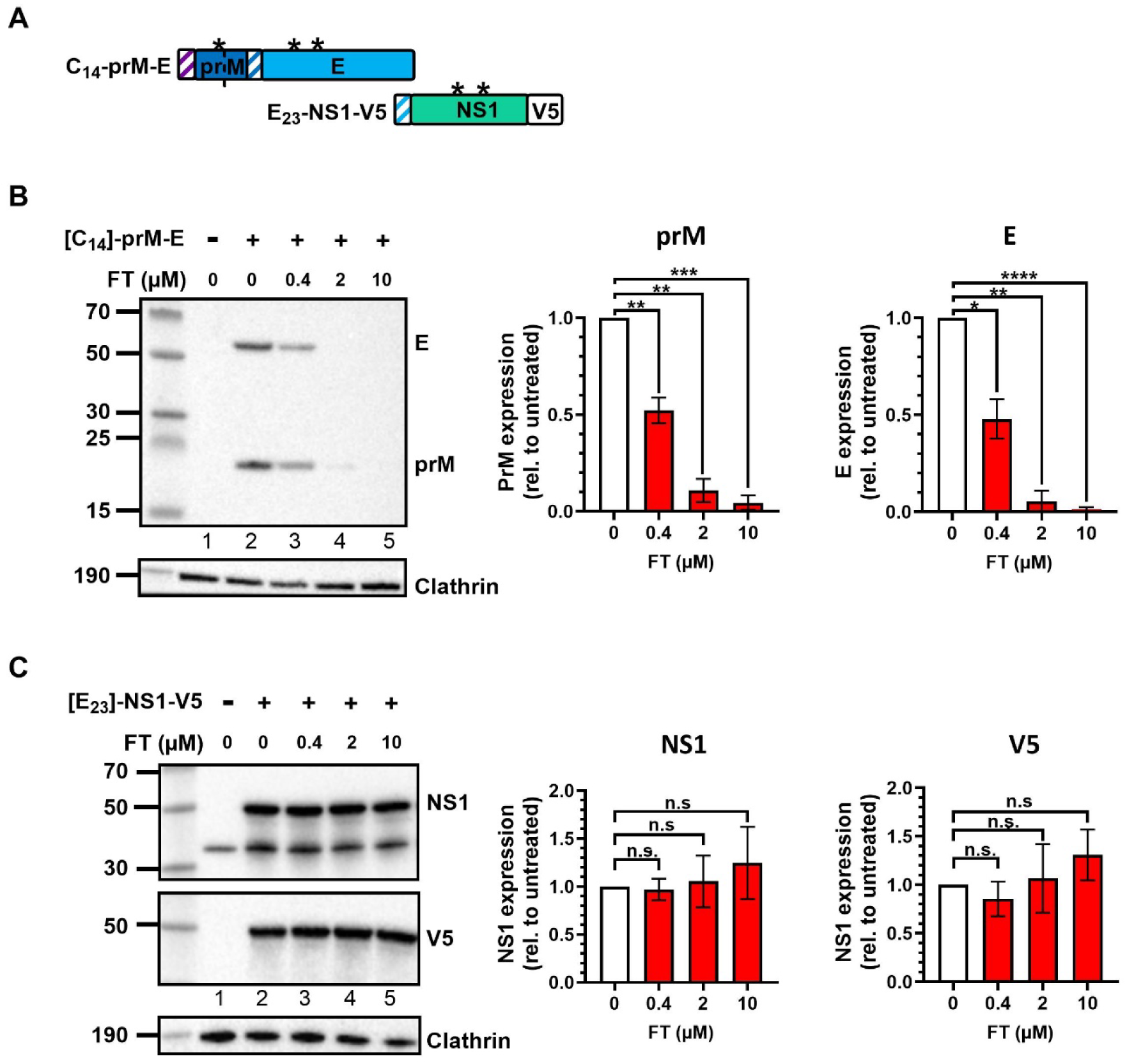
Flavitransin inhibits DENV2 prM-E protein expression in transiently transfected HEK293T cells. **A**, Schematic representation of the plasmid constructs expressing DENV2 C_14_-prM-E and E_23_-NS1-V5. N-linked glycosylations are indicated by an asterisk. **B,** HEK293T cells were transiently transfected with the C_14_-prM-E construct and treated with FT for 18 h. Cells were lysed in NP-40 buffer and analyzed by immunoblotting with an anti-E or an anti-prM antibody. For the cell loading control, an anti-clathrin antibody was used. A representative immunoblot is shown on the left. Bar graphs represent relative prM and E levels from clathrin-corrected samples, normalized to untreated transfected controls. Bars are mean ± SD, n=3. *P< 0.05, **P<0.01, ***P<0.001, ****P<0.0001, compared to untreated transfection control (two-tailed/multiple unpaired t test with Welch’s correction). Note that the processing of prM into pr and M by furin cleavage was not successful in HEK293T cells. **C**, Same as in (B) but with the E_23_-NS1-V5 construct and with anti-NS1 or anti-V5 antibodies. n.s. = P>0.05.

The second plasmid encoded for the non-structural NS1 protein that was modified with an additional V5 tag at its C-terminus. NS1 carries the hydrophobic C-terminal end of the envelope protein at its N-terminus that serves as a cleavable SP (**Fig. 1A** and **Fig. 2A**). In sharp contrast to prM and E, expression of NS1 protein was not suppressed by FT treatment at concentrations up to 10 µM, as determined by immunoblotting of cell lysates with an anti-NS1 or an anti-V5 antibody (**Fig. 2C**, lane 3-5). This was confirmed at the secretion level of NS1 as evidenced by comparable protein levels of NS1 in the cell culture medium in the absence or presence of FT or FT_IN_ (**Suppl. Fig. 4**). Altogether, these results indicate that the antiviral effect of FT is related to a direct inhibitory effect on the expression of the structural proteins of the viral envelope.

### Flavitransin inhibits the translocation of the DENV prM protein

Next, the direct effect of FT on translation and/or translocation of the separate proteins prM and E was further established by means of an in-house *in vitro* cell-free rabbit reticulocyte lysate assay, supplemented with ovine pancreatic rough microsomes (RM) that represent the ER (**Fig. 3A**)[47]. Translation of the prM transcript was not affected by FT (**Fig. 3B**, lane 2, black arrowhead). However, in the presence of RM (lanes 3-12), translocation of prM into the ER lumen, as evidenced by the appearance of glycosylated species with a higher molecular mass (lane 3, asterisk), was significantly inhibited by a 15 µM FT treatment (lane 4) but not by the same concentration of the control compound FT_IN_ (lane 7). Translocation inhibition by FT was further confirmed through proteinase K analysis of the samples, showing a clear impact on the amount of rescued glycosylated protein in the protected ER lumen for the FT sample (**Fig. 3B**, lane 9) as compared to the untreated control (lane 8).

**Figure 3.**
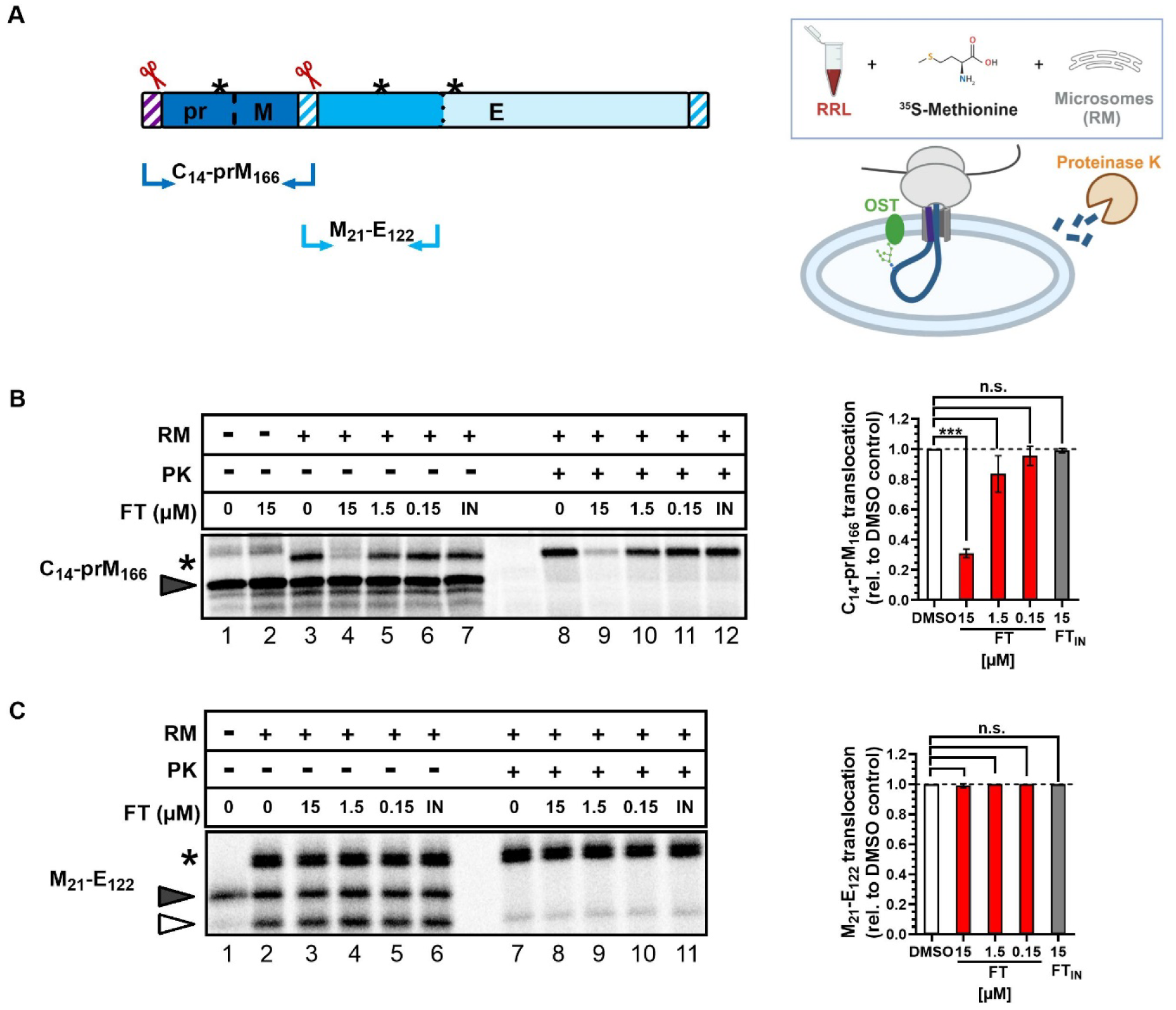
Flavitransin inhibits Sec61-directed co-translational translocation of DENV2 prM protein but not E protein *in vitro*. **A**, Schematic representation of the DENV prM and E transcripts for *in vitro* translation/translocation assay in which the transcripts are translated in rabbit reticulocyte lysate (RRL) supplemented with ovine rough microsomes (RM) and [^35^S]-labeled methionine. Only fully translocated proteins in the ER lumen are protected from proteinase K (PK) treatment. Red scissors indicate signal peptidase cleavage sites and the asterisks represent the N-linked glycosylation sites. Cartoon created with BioRender (2025); www.bioRender.com. **B**, Transcripts encoding DENV2 C_14_-prM_166_ were translated and/or translocated in the assay described in (A). Samples were left untreated or were treated with the indicated concentrations of FT or with 15 µM of FT_IN_. Half of the translated material of each sample was additionally treated with PK. Samples were separated by SDS-PAGE and analyzed by autoradiography. A representative autoradiogram is shown on the left. Graph on the right shows the relative translocation as normalized to the DMSO control from 2 independent experiments. Bars represent mean +/- SD; n=2. ***P<0.001; n.s. = P>0.05 for multiple unpaired t tests with Welch’s correction. Solid arrowhead: unprocessed nascent protein; asterisk: glycosylated (translocated) mature protein. **C**, Same as in (B) but for the M_21_-E_122_ construct. Open arrowhead: SP-cleaved protein that is not being glycosylated. Only a small fraction of these species are translocated into the ER lumen and protected from PK.

Because of the large size of the full length E protein (495 AA), we opted to work with a truncated variant of 143 residues, in order to increase the success rate of its translation in a cell-free system (**Fig. 3A**). *In vitro* translocation of the E protein results in the generation of 2 additional species to the translated pre-protein: a non-glycosylated fraction that is cleaved by the signal peptidase (**Fig. 3C**, lane 2; open arrowhead), and a glycosylated fraction with a higher molecular mass (lane 2, asterisk). In contrast to the effect on prM and the *in cellulo* findings of transfected HEK293T cells (**Fig. 2**), translocation of the isolated E protein alone was not affected by FT (lanes 3-5), nor by the control compound FT_IN_ (lane 6). These results indicate that FT selectively inhibits the translocation of the prM segment of the DENV polyprotein across the ER membrane.

### Flavitransin diverts the DENV polyprotein to the proteasome for degradation

We hypothesized that the inhibitory effect of FT on the translocation of the prM subunit halts the subsequent insertion of the whole viral polyprotein into the ER membrane, and initiates the rerouting of the pre-protein to the cytosol where it gets degraded by the proteasome. To track the different subunits of the viral envelope proteins (i.e., pr, M and E), a construct was designed in which a different protein tag was introduced in each of the subunits for identification purposes (**Fig. 4A**). Cells were transfected and subjected to a combinational treatment of FT (or FT_IN_) and the proteasome inhibitor MG132. In the absence of compounds, the tagged prM (V5 and FLAG) and E (MYC) proteins were successfully expressed and glycosylated, as confirmed by endoglycosidase H (Endo-H) treatment (**Fig. 4B** and **C**), indicating a correct topology of the modified construct. Interestingly, a 75 kDa precursor (prM/E) could be rescued by the addition of both FT and MG 132 (**Fig. 4B** and **Suppl. Fig. 5A**, lane 8-10, arrow), as evidenced from the detection of the 3 protein tags (V5, FLAG and MYC) that span the whole pre-protein, and by additional V5 pull-down experiments (**Fig. 4D**). Of note, this precursor could already be detected by single FT treatment (**Fig. 4B**, **Fig. 4D** lane 3 and **Suppl. Fig. 5A** lane 5). Moreover, Endo-H treatment of the MG132 samples indicated that the 75 kDa precursor remains in a non-glycosylated form (**Suppl. Fig. 5B**, arrow). All together, these data suggest that the mis-targeted viral protein that does not reach the ER lumen under FT pressure is re-routed to the cytosol for proteasomal degradation.

**Figure 4.**
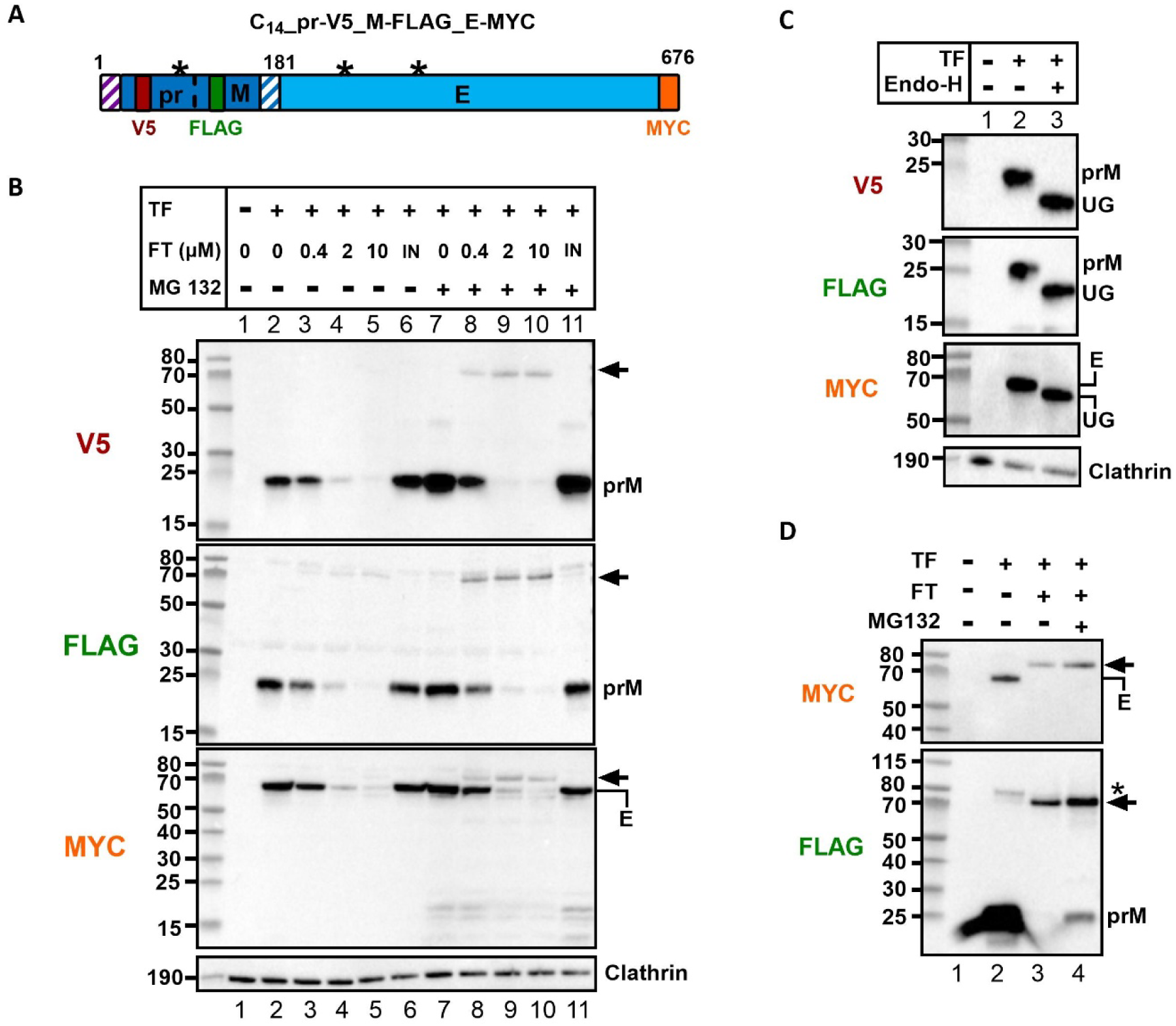
Flavitransin diverts the untranslocated C_14_-prM-E precursor to the proteasomal degradation pathway. **A**, Schematic representation of the modified DENV2 C_14_-prM-E construct with a V5 tag in the pr protein, a FLAG tag in the M protein and a MYC tag in the E protein (tags are represented by colored boxes). Transmembrane domains are indicated by shaded boxes. N-glycosylation sites are indicated by an asterisk. **B**, CHO-K1 cells were transiently transfected (TF) with the construct from (A) and treated with FT, FT_IN_ (10 µM) and/or the proteasome inhibitor MG132 (200 nM). Compound treatment was started at 6 h post transfection for 18 h and cells were lysed in NP-40 buffer and analyzed by immunoblotting. For the cell loading control, an anti-clathrin antibody was used. The uncleaved precursor (prM-E) is indicated by a black arrow. One representative experiment out of three is shown. **C**, Cell lysate of control samples from (B) was incubated with Endo-H for removal of N-glycosylation and subjected to immunoblotting like in (B). UG: unglycosylated. **D**, V5 pull down. Samples from (B) were incubated with anti-V5 beads for pull down of the V5-tagged protein species and immunostained with anti-FLAG and anti-MYC antibodies. Black arrow indicates the uncleaved precursor that is already detectable in the FT sample without MG132. Note that some processed E protein is co-immunoprecipitated with prM as viral envelope protein complexes (MYC detection). In the control sample, an additional species with a higher molecular weight is detected (*), presumably a glycosylated precursor protein (∼ 80kDa) that was not properly cleaved into the prM and E subunits.

### The C-terminus of DENV capsid protein determines sensitivity to Flavitransin

Next, we zoom in on the region of the prM subunit that confers sensitivity to FT. The viral envelope glycoprotein (G) of Vesicular Stomatitis Virus (VSV) was selected as our negative control as it proved to be resistant to FT (data not shown). VSV-G, a type I protein with a N-terminal SP and a C-terminal TMD as membrane anchor, was elongated with a GFP-P2A-BFP sequence (**Fig. 5A**) for downstream flow cytometric analysis, similar to other modified proteins in earlier studies [48, 49]. As anticipated, VSV-G-GFP expression in transiently transfected HEK293T cells was not affected by FT (**Fig. 5A**; grey curve). Additional constructs were designed in which the N-terminal part of VSV-G was exchanged by the corresponding region of the DENV proteins prM, E and NS1. Specifically, in addition to the SP-like targeting TMD sequence (i.e., C_14_, M_21_ and E_23_), 62 residues of the mature protein of each DENV subunit were also introduced in the VSV-G backbone (**Fig. 5A**) to preserve translocation features of the respective wild-type DENV protein [50]. Interestingly, the expression of the prM-G chimeric protein was profoundly suppressed by FT treatment in a concentration-dependent manner (IC_50_ = 283 nM, **Fig. 5A**; purple curve), in line with our data with full-length wild-type prM protein in infected (**Suppl. Fig. 2**) and transfected (**Figs. 2** and **4**) cells and in cell-free *in vitro* translocation assays (**Fig. 3**). In contrast, FT had only a minor effect on the E-G and NS1-G chimeric constructs (IC_50_ > 10 µM, **Fig. 5A**; blue curves). As expected, all constructs showed resistance to the treatment with the control compound FT_IN_ (**Suppl. Fig. 6A**). Further constriction of the DENV protein part to the SP-like targeting sequence only in the DENV-VSV chimaeras revealed that the C_14_ sequence is the main determinant to FT sensitivity. Exchanging the SP of VSV-G by the C_14_ sequence of prM resulted in a VSV-G protein that became highly regulated by FT (IC_50_ = 44 nM, **Fig. 5B**), whereas FT had no down-modulating effect on the M_21_ targeting sequence (IC_50_ > 10 µM, **Fig. 5B**), but partially suppressed the expression of the G chimaera with the E_23_ targeting sequence (IC_50_ = 1.91 µM, **Fig. 5B**).

**Figure 5.**
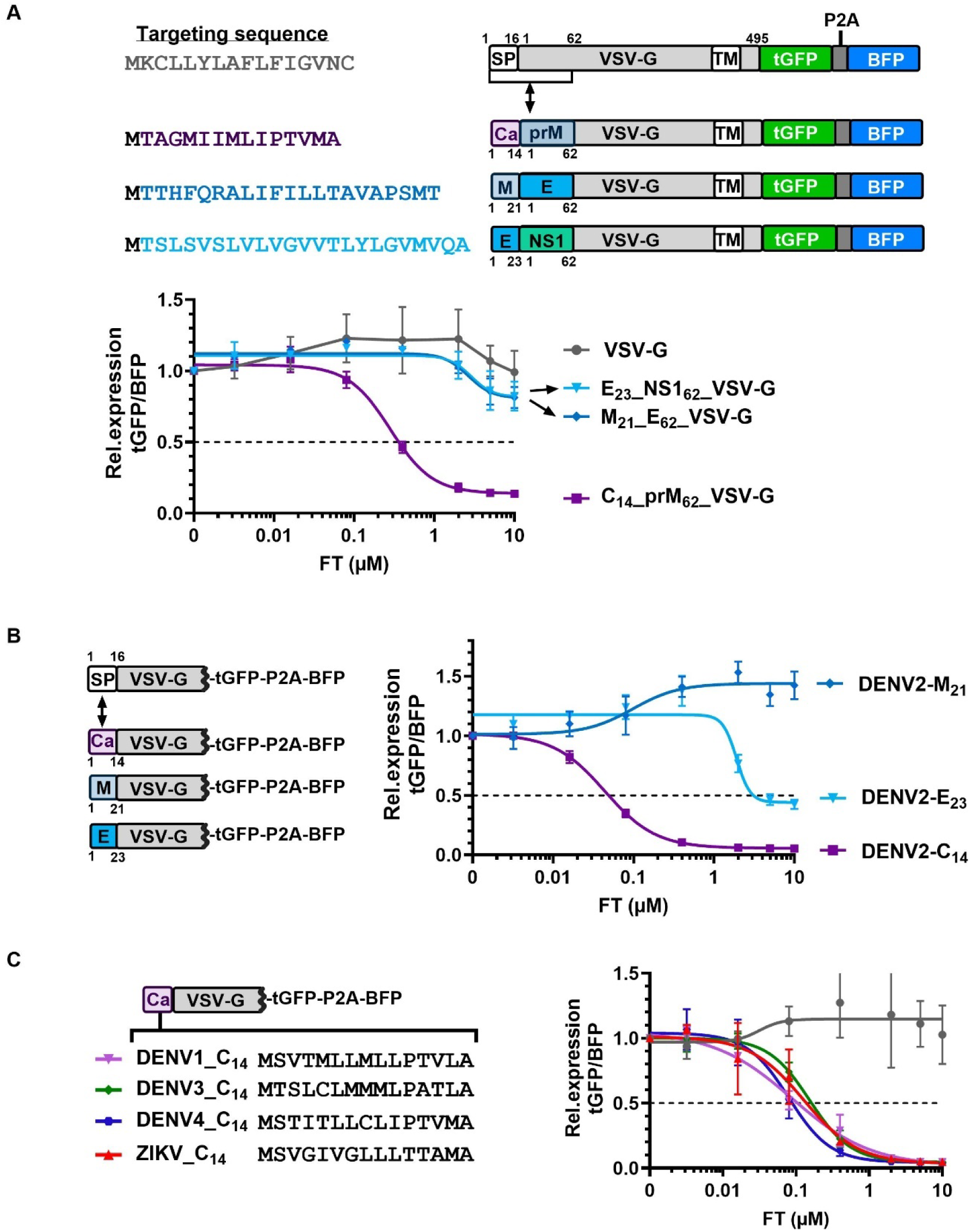
Flavitransin selectively targets the membrane-spanning domain of the capsid subunit in the flaviviral polyprotein. **A**, Schematic representation of the constructs used for transfection and flow cytometry. In the VSV-G backbone with tGFP-P2A-BFP sequence, the SP and 62 AA of the mature G protein are exchanged by the corresponding signal peptide-like targeting sequence and 62 AA of the mature DENV subunit. The AA sequence of each targeting sequence is shown. Note that an extra methionine (black) is added at the N-terminus of each DENV targeting sequence for translation initiation. HEK293T cells were transiently transfected, treated with FT for 18 h, and analyzed by flow cytometry for tGFP and BFP expression levels. Graph shows the four parameter concentration-response curves of FT for the different constructs for tGFP expression (mean fluorescent intensity). Relative protein expression was calculated by the ratio tGFP:BFP, normalized to untreated transfection control for each construct (set at 1.0). Values are mean ± SD; n=3. Note that the two blue curves coincide. **B**, Same as in (A), but for constructs in which the SP only of G was exchanged by the corresponding signal peptide-like targeting sequences (C_14_, M_21_ or E_23_) of the different DENV subunits. Graph shows mean ± SD; n=3. **C**, Same as in (B) but for the C_14_ signal peptide-like targeting sequences of the other serotypes of DENV and of ZIKV. The AA sequence of each C_14_ SP is shown. Note that an extra methionine is added at the N-terminus of each flaviviral targeting sequence for translation initiation. Graph shows mean ± SD; n=3. The grey curve represents the VSV-G-tGFP-P2A-BFP control with the SP of WT VSV-G.

These data show that, in sharp contrast to prM, all of the tested DENV2 proteins and the VSV-G protein are not or nearly not affected by FT, suggesting a certain client selectivity for this translocation inhibitor. To further explore this, endogenous cell surface expression of several type I TM receptors was determined on MT-4 cells in the absence or presence of FT. Little impact of FT was recorded for most receptors (**Suppl. Fig. 6B**), except for CD4 for which a profound suppression was seen as reported earlier [51]. However, the impact of FT on the DENV2 prM protein was slightly bigger as compared to CD4.

### Flavitransin exerts broad anti-flaviviral activity

Next, we explored the antiviral activity of FT in additional relevant cell types and against different variants of DENV. In monocyte-derived dendritic cells (MDDC), one of the first target cells of flaviviruses in the human skin shortly after a mosquito bite, DENV2 infection was strongly suppressed by FT, but not by its inactive analogue FT_IN_ (**Table 1** and **Suppl. Fig. 7A**). Likewise, in human B-cell lymphoma Raji cells stably transfected with the DENV entry receptor CD209 (better known as Dendritic Cell-Specific Intercellular adhesion molecule-3-Grabbing Non-integrin; DC-SIGN), DENV2 infection was concentration dependently inhibited by FT (**Table 1** and **Suppl. Fig. 7B**), whereas the cellular expression of CD209 remained unaffected by FT. Furthermore, FT preserved activity, although to a lesser extent, in the DENV susceptible C6/36 cells derived from larva of *S. albopictus*, the principal vector of DENV transmission (**Table 1** and **Suppl. Fig. 7C,D**). Interestingly, in a cell-free insect lysate assay (from *Spodoptera frugiperda Sf12*), FT also inhibited the translocation of the DENV prM protein as evidenced by the almost complete absence of the glycosylated (hence translocated) protein (**Suppl. Fig. 7E** lane 2 and 5). This supports that the translocation inhibitory effect of FT is preserved among human and insect cells, explaining the broad-spectrum antiviral effect in both host and vector (**Table 1**).

Importantly, comparable sub-micromolar activity of FT was recorded against a panel of laboratory-adapted strains and clinical isolates belonging to the four DENV serotypes (**Table 1**), demonstrating a pan-DENV-serotype effect of FT. Furthermore, FT exerted antiviral activity against two other members of the flavivirus genus. FT prevented ZIKV infection at sub-micromolar concentrations comparable to that of DENV4 (**Table 1** and **Suppl. Fig. 3B**), and exerted a substantial antiviral effect on YFV infection (**Table 1**). As expected, FT_IN_ remained inactive against all the tested flaviviral strains (data not shown). Altogether, these data demonstrate a consistent antiviral potency of FT in different cell types, originated from different species, and a pan-flavivirus antiviral activity.

To investigate this pan-flaviviral potency of FT in more detail, VSV-G mutants were generated in which the SP of G was exchanged for the SP-like targeting C_14_ sequence of prM of the other DENV serotypes and ZIKV (**Fig. 5C).** Remarkably, whereas the C_14_ AA sequence for the 4 serotypes and ZIKV showed quite some variability between one another, the expression of all the C_14_-VSV-G constructs was concentration-dependently suppressed by FT in a similar way (IC_50_ values ∼ 100 nM) (**Fig. 5C**), suggesting that some secondary characteristics of the SP and/or Sec61 translocon are at play in FT sensitivity.

### Flavitransin-desensitized cells express resistance mutations in Sec61α

By analogy with the CADA derivative CK147 [45], we next explored if FT interacts with the Sec61 translocon. Therefore, HCT116 cells were cultivated in high concentrations of FT to generate escape mutants that survive the (sub-)toxic dose. Because of a rather limited cytotoxic (but more cytostatic) effect of FT on cells (**Fig. 1D** and **Suppl. Fig. 8A**), we added extra pressure to the cells by cultivating them in suspension at very low density during the initiation of FT exposure (**Fig. 6A**). Cells that survived this FT treatment were clonally expanded and subjected to further in-depth phenotypic and genotypic analysis. From two independent attempts, we selected five HCT116 clones that became desensitized to FT, as determined by the cell proliferation profile at high FT concentrations (CC_50_ values > 50 µM; **Fig. 6B**). Next, the sequence of the Sec61α subunit was analyzed in the resistant clones and aligned to that of WT HCT116 cells. We identified four different heterozygous mutations in Sec61α across the resistant clones, i.e., A70V, E78A, E232K and L449F, that arose from three clones with single mutations and two dual mutants (**Fig. 6B**).

**Figure 6.**
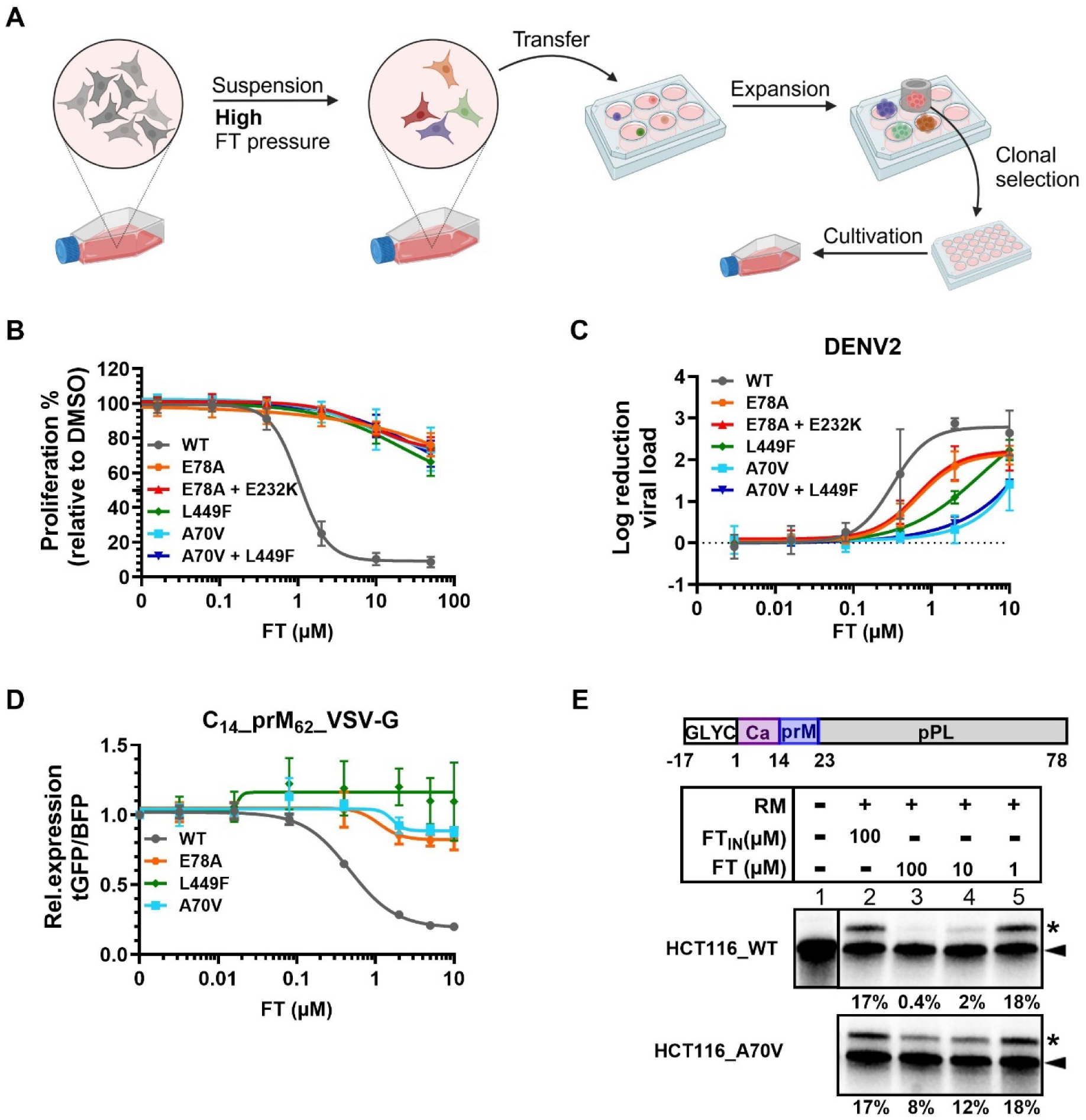
Generation and analysis of flavitransin-desensitized HCT116 cells. **A**, Schematic outline of the protocol used to select for FT-desensitized cells. HCT116 cells were seeded at low density in suspension in the presence of FT (25 µM). After 9 days, surviving cells were subcloned, expanded and genotyped. Cartoon created with BioRender (2025); www.bioRender.com. **B**, Four-parameter concentration-response curves of FT for the proliferation of WT and FT-resistant HCT116 cells. After 72 h of FT incubation, cell proliferation was quantified by measuring resazurin:resorufin conversion. For each cell clone, data are normalized to the DMSO control. Values are mean ± SD; n=3. **C,** WT and FT-resistant HCT116 cells were infected with DENV2 (MOI 1) and incubated for 5 days in the presence of various FT concentrations. Viral copy number in the supernatant was quantified by RT-qPCR. Graph shows the four-parameter concentration-response curves of FT for viral replication (log_10_ reduction in viral load compared to untreated infected controls). Values are mean± SD; n=3. **D**, WT and FT-resistant HCT116 cells were transiently transfected with a plasmid encoding for DENV2 C_14_-prM_62_-VSV-G protein (same as in Fig. 5A) and treated with FT for 18h. GFP and BFP expression (mean fluorescent intensity) was quantified by flow cytometry and normalized to untreated transfection control. Graph shows the four-parameter concentration-response curves of FT for the tGFP:BFP ratio. Values are mean ± SD; n = 3. See also Suppl. Fig. 8B,C. **E**, WT and Sec61α A70V mutant HCT116 cells were semi-permeabilized with digitonin and used as RM source in an *in vitro* translation/translocation assay. Transcripts from a N-Glyc-C_14_-prM_8_-pPL construct were translated and translocated in rabbit reticulocyte lysate with [^35^S] methionine, FT or FT_IN_ (100 µM), and extracted membranes from semi-permeabilized cells. The pPL construct contains the C14 sequence of DENV that is N-terminally elongated (17 residues) with a N-glycosylation sequon as previously described [43]. Samples were separated by SDS-PAGE and analyzed by autoradiography. The percentage of translocated (glycosylated) protein (relative to unprocessed precursor) is marked below each lane. Translation in absence of membranes produces the unprocessed nascent protein (solid arrowhead). N-glycosylated species are indicated by an asterisk. One out of three representative experiments is shown. See also Suppl. Fig. 8D.

To verify FT resistance, we analyzed the impact of the Sec61α mutations on the antiviral potency of FT against live DENV2. Fortunately, HCT116 cells proved to be permissive for DENV infection and could be employed for viral replication assays. As shown in **Fig. 6C**, WT HCT116 cells responded well to FT which inhibited DENV2 replication concentration-dependently (IC_50_ value of 319 nM and a 3 log_10_ reduction in viral load with 2 µM FT as compared to the untreated infected control). FT treatment of infected HCT116 cells with mutant Sec61α resulted in a clear reduced antiviral effect on DENV2 replication (**Fig. 6C**). Both single E78A and double E78A + E232K mutant cells responded to FT in a similar way and showed decreased sensitivity to FT in suppressing DENV2 replication (2 log_10_ reduction in viral load with 2 µM FT) as compared to the WT cells (3 log_10_ reduction; **Fig. 6C**). The identical FT concentration-response curves for the E78A and the E78A + E232K mutant cells, suggest that E232K is a potential compensatory mutation. The strongest FT-resistance was recorded for the A70V mutant, for which only a 0.3 log_10_ reduction in viral load was obtained with a 2 µM FT treatment. Although the L449F amino acid substitution in Sec61α had a clear impact on FT sensitivity as a single mutation (1 log_10_ reduction in viral load with 2 µM FT), the combination with A70V did not result in an additive effect, suggesting a dominant role of this A70V amino acid substitution in FT resistance (**Fig. 6C**). In accordance with the effect on live WT virus (**Fig. 6C**), the Sec61α mutations had a direct impact on DENV2 prM escape from FT suppression as assessed by C_14_-prM_62_-VSV-G reporter protein expression. In transiently transfected WT HCT116 cells, prM expression was concentration-dependently down-modulated by FT (**Fig. 6D**; grey curve; IC_50_ = 470 nM), in a comparable manner as was seen in transfected HEK293T cells (**Fig. 5A**). However, in the Sec61α mutant cells, C_14_-prM_62_-VSV-G expression was no longer affected by FT (**Fig. 6D** and **Suppl. Fig. 8B, C**), validating FT resistance of the Sec61α mutants on DENV2 prM protein translocation. Furthermore, in a cell-free context with ER membranes extracted from semi-permeabilized WT and mutant HCT116 cells, import of a prM-prolactin reporter into the ER lumen was clearly less suppressed by FT treatment in the A70V mutant as compared to WT Sec61α as evidenced by the fraction of glycosylated (= translocated) species (**Fig. 6E** and **Suppl. Fig. 8D**). A similar reduced effect of FT on the expression of the prM protein in the A70V mutant was detected in transiently transfected HCT116 cells by immunoblotting (**Suppl. Fig. 8E**).

### Flavitransin has a high barrier to viral resistance

Finally, as viruses are known to be masters in generating resistant mutants that can escape antiviral pressure, we determined the resistance barrier to FT. Specifically, WT DENV2 virus was passaged in Huh7 cells in presence of gradually increasing sub-inhibitory concentrations of FT, starting from a 0.1 µM dose (**Fig. 7A**). Resistance selection to FT was performed in two independent experimental procedures (indicated as FT_1_ and FT_2_). In addition, virus was passaged in cells exposed to the inactive FT_IN_ compound and in cells that received only culture medium (indicated as Control). More than 75 passages were needed to select virus that could replicate in a 14-fold elevated FT dose (1.35 µM), whereas this concentration of FT completely prevented replication of the original WT virus (**Fig. 1E**). As shown in **Fig. 7B**, the selected FT-exposed DENV strains showed reduced sensitivity to FT but remained fully blocked in their replication at a high FT concentration (2 µM). Passaging of the virus in the presence of FT_IN_ or control medium also resulted in a (slightly) reduced sensitivity to FT, presumably by introduction of random mutations throughout the viral genome that affect viral fitness (**Fig. 7B, C**). Whole-genome sequencing of the cultivated DENV strains collected at end-point (passage 75) revealed several AA substitutions throughout the viral genome as compared to the initial virus control (**Fig. 7C**). In the FT-selected virus strains, several non-synonymous mutations were detected in the viral genes that resulted in unique AA substitutions in the Ca (N79K; A102V), the M (L4F; I53T) and the E (P80S; H346Y; V354A) proteins, that were present in both independent FT-exposed strains, but that were not selected in the control viruses that underwent the same passaging in FT_IN_ compound or culture medium (**Fig. 7C**). Interestingly, we identified a A102V mutation in the capsid protein that was located in the C_14_ targeting sequence of prM (mutation detected in 99 % of all viral reads).

**Figure 7.**
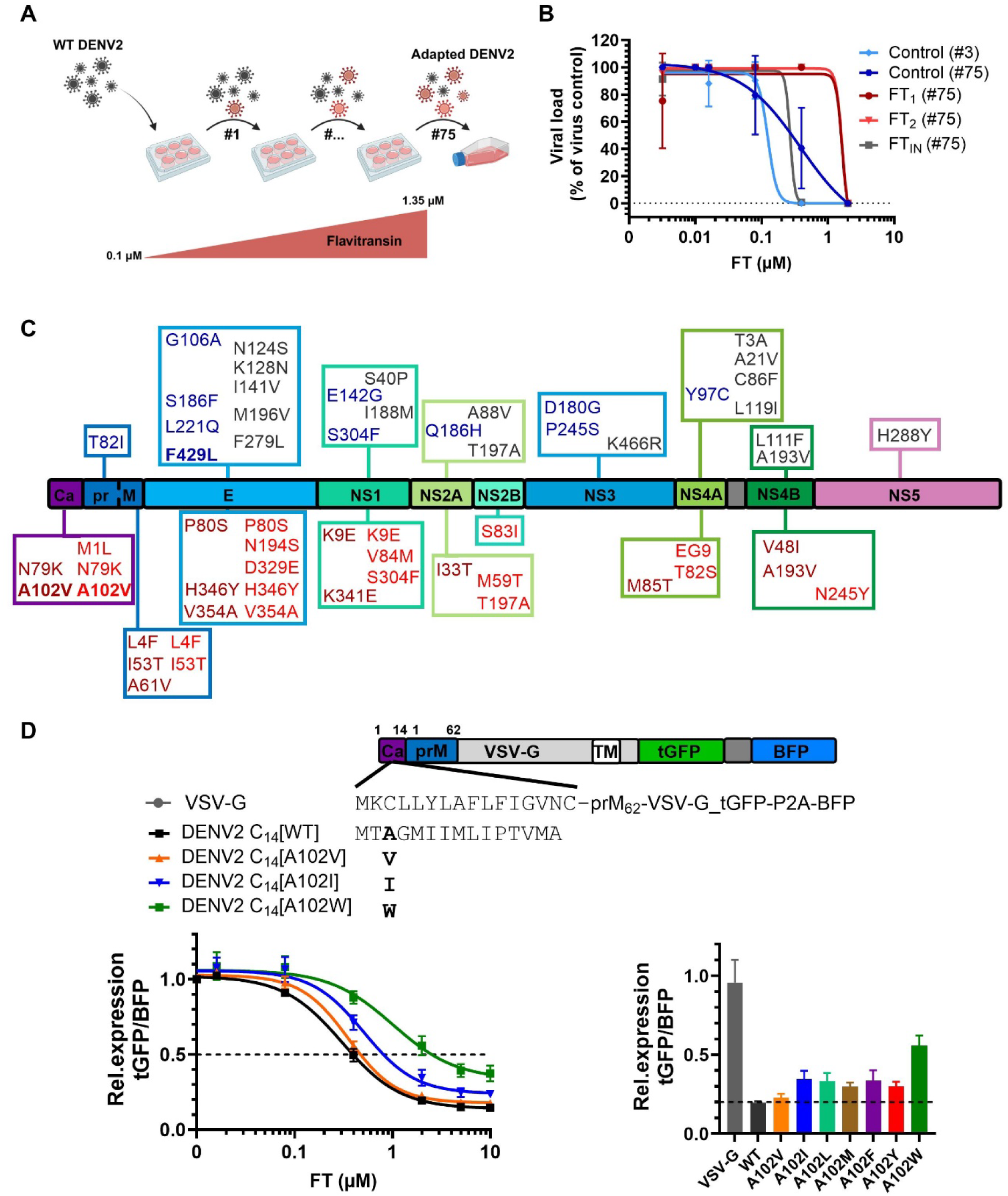
Generation and analysis of flavitransin-desensitized DENV2 virus strains. **A**, DENV2 was passaged on Huh7 cells under FT pressure. Compound concentration was gradually increased when viral cytopathic effect on cells was detected. After 75 passages, virus was cultivated in the absence of compounds to generate virus stocks for downstream analysis. At passage 50 and 75, virus was collected and genotyped by nanopore sequencing (see panel C). FT resistance was selected in 2 independent cell cultures. As controls, a culture with FT_IN_ or with control medium was included that underwent the same passaging procedure. Cartoon created with BioRender (2025); www.bioRender.com. **B**, At end-point, virus stocks were evaluated in Huh7 cells for sensitivity to FT. A virus stock from a low passage (# 3; light blue) was included as reference. Cells were infected with the different virus stocks (MOI ∼ 0.5) and treated with increasing concentrations of FT. After 4-5 days post infection, virus yield in the supernatant was quantified by RT-qPCR detection of the viral 3’UTR. Graph shows the four parameter concentration-response curves of FT for virus release from infected cells. Values (mean ± SD, from two independent experiments; n=2) represent the viral load in FT-treated cells as normalized to the viral copy number of each untreated virus control. **C**, Summary of the detected AA mutations in the different viral clones as compared to the initial DENV2 control strain (# 3). The numbers in each colored box refer to the residues of the respective viral subunit. The mutations in the medium control (dark blue) and FT_IN_ (grey) are shown at the top; the two FT-resistant viruses (dark and light red) are shown at the bottom. **D**, Schematic representation of the hydrophobic mutants of the DENV2 C_14_ SPs. HEK293T cells were transiently transfected with the VSV-G-tGFP-P2A-BFP constructs and treated with FT for 18 h. GFP and BFP fluorescence (mean fluorescent intensity) was quantified by flow cytometry and relative tGFP:BFP expression ratio (normalized to untreated transfection control) was determined. Graph on the left shows the four parameter concentration response curves of FT for VSV expression for different C14 mutants from three independent experiments. Values are mean ± SD; n=3. Graph at the right shows the relative tGFP:BFP expression ratios at 2 µM FT for additional hydrophobic mutants (A102V/I/L/M/F/Y/W). Values are mean ± SD; n=3.

To directly assess the effect of the A102V mutation in the DENV protein on FT resistance, we introduced the A102V amino acid exchange in our C_14_-prM-VSV-G reporter protein for transfection in HEK293T cells (**Fig. 7D**). The impact of this A102V mutation in prM for FT resistance was very little, with only minor (nearly detectable) reduced sensitivity to FT as compared to the WT reporter (**Fig. 7D**). As this exchange of alanine (capsid residue 102) by a more hydrophobic valine might enhance the general hydrophobicity of the C14 sequence, we further explored the impact of substituting alanine at this position with more hydrophobic amino acids (such as, I, L, M, F, Y and W). Thus, additional hydrophobic C14 mutants were generated as shown in **Fig. 7D**. Interestingly, substituting alanine by an isoleucine residue had a clear effect on FT sensitivity, which was most pronounced when exchanging alanine by a tryptophan residue.

## DISCUSSION

DENV remains a significant public health concern as its mosquito vector continues to spread to new areas worldwide. Current treatment options are limited to supportive care, including fluid administration and antipyretic therapy, since no specific antivirals for DENV (or other flavivirus infections) are approved. Therefore, there is an urgent need for effective pan-serotype DENV antivirals that can reduce viral load during infection and mitigate DENV-associated clinical manifestations. The most desirable inhibitor would possess pan-flavivirus activity, considering the risk of sudden (re)- emergence of flaviviruses, such as the ZIKV outbreak in 2015 [52].

In this study we explored the antiviral potency of the CADA derivative FT against DENV and related flaviviruses and observed a potent and consistent antiviral effect across the 4 serotypes of DENV, ZIKV and YFV. Given the structural resemblance of FT with the earlier reported Sec61α inhibitors CADA and CK147 [43, 45], we anticipated that this strong antiviral effect of FT was related to inhibition of the Sec61-dependent co-translational translocation process at the ER. In the context of virus propagation, preventing protein import into the ER by Sec61α inhibitors can affect the expression of either cellular receptors used by the virus to attach to and enter the host cell, as has been described for the human CD4 receptor of HIV and CADA [42, 43], or viral envelope proteins crucial for host cell attachment and membrane fusion as has recently been reported for mycolactone [41]. From our time-of-drug-addition data (**Fig. 1H**), we could exclude a direct inhibitory effect of FT on cell surface proteins that serve as viral entry receptors, given that we could delay FT treatment by 8 h and still completely block viral replication. However, the broad-spectrum (pan-flaviviral) antiviral effect of FT would rather suggest a common cellular factor as drug target, for example the cellular receptor DC-SIGN used both by DENV and ZIKV [53, 54], but the expression of this type II receptor was not affected by FT (**Suppl. 7B**).

Genome-wide screens identified the ER as a key organelle throughout the viral life cycle of DENV [9–12, 14]. The Sec61 translocon and several of its accessory components (such as signal recognition particle complex, TRAP subunits, and Sec62/63) were previously reported as host-dependency factors for flavivirus replication, and Sec61 inhibition was shown to block DENV and ZIKV infection [13, 39, 41]. A unique feature of flaviviruses is that their viral RNA genome is translated into a single, large polytopic protein, containing 18 membrane-spanning domains that, similar to cellular multi-pass transmembrane proteins, require insertion in the ER membrane (**Fig. 1A**). As a consequence, all of the viral protein subunits (both the structural and the cytosolic positioned non-structural proteins) rely on the ER for their expression. The DENV polyprotein is a rather atypical (and complex) multi-pass transmembrane protein in cells as it contains four internal signal peptidase cleavage sites, comparable to cleavable SPs of type I transmembrane proteins. Through a set of cell-transfection experiments and complimentary cell-free in vitro translation/translocation assays we could irrefutably demonstrate that only one of the viral SP-like TMDs confers sensitivity towards FT, i.e., the first (N-terminal) hydrophobic region that is being translated and serves as the main Sec61 targeting sequence of the viral nascent chain. In contrast, the more downstream SP-like TMDs of the E and NS1 protein showed full resistance to FT. Comparison of the different targeting sequences (**Fig. 5A**) revealed a clear difference in SP length (14 residues vs 21 and 23), but not in hydrophobicity (based on the Kyte-Doolittle hydrophobicity scale), 2 parameters that can contribute to SP efficiency [48, 55–58]. Considering the function of the SP-like TMDs of E and NS1 as membrane anchors of prM and E, respectively, for expression at the cell surface and anchoring in the viral envelope, these TMDs are supposed to be sufficiently hydrophobic. The unexpected co-IP of the E protein with prM (**Fig. 4D**, lane 2) demonstrates a native expression of these viral proteins in transfected cells with a clear interaction between both envelope proteins, which is in line with the reported protective role of prM for E stability [59]. The fourth SP-like TMD, i. e., the 2K peptide between NS4A and NS4B protein was not assessed in our study, but the biogenesis of NS4A and NS4B was previously shown to be dependent on the EMC [17, 18].

From our experiments with the proteasome inhibitor MG132 we concluded that in the presence of FT the nascent chain remains in its pre-form and gets degraded by the proteasome. The rescue of the pre-protein was most clear from the combinational treatment of FT with a proteasome inhibitor, but already detectable (although at very low amounts) with the Sec61 inhibitor only, as has also been observed for some CADA clients [60]. The absence of N-glycosylation of this precursor indicates that the nascent protein chain is certainly not reaching the ER lumen. As both prM and E are fully degradable by proteinase K as verified in our cell-free *in vitro* translocation assay, it is unlikely that the prM-E precursor would survive in the cytosol without blockage of the proteasome. This suggests that some of the nascent prM-E protein of a stalled ribosome might be trapped by FT in the protected vestibule of ribosome exit tunnel/translocon channel that escapes protein degradation.

The selectivity of FT for the C14 sequence within the flavivirus polyprotein is a unique observation and interesting, and might explain the strong impact of FT in aborting the vulnerable translocation initiation at Sec61 for successful ER import. However, it somehow contradicts the pan-serotype and broad-flavivirus activity of FT given the considerable variation in the C14 targeting sequences of the different virus strains (**Fig. 5C**). This again argues for the importance of secondary or other intrinsic features of SPs besides the primary AA sequence that contribute to sensitivity to Sec61 inhibitors, as has also been proposed for the cotransins [57]. In line with this, the observed A102V mutation in the selected FT-desensitized DENV strain might as single AA exchange not be sufficient for FT escape, given that both WT DENV serotype 1 and WT ZIKV carry a valine residue at the same position (**Fig. 5C**) but are fully sensitive to FT. This is in favor of a high barrier to resistance, suggesting that multiple mutations in this C14 sequence are required to escape drug pressure, but without turning this SP-like sequence into a non-functional targeting signal. Nevertheless, one escape route for the virus could be the selection of mutant strains with enhanced hydrophobicity in the C14 targeting sequence (as we demonstrated for the tryptophan mutant; **Fig. 7D**), which might result in a ‘stronger’ SP. However, this would demand multiple simultaneous non-synonymous substitutions in the viral RNA genome that result in a AA mutation (the A102V mutation only required one nucleotide GCA to GTA substitution). The nearly detectable FT resistance of a single A102V mutation in our reporter construct should be seen in the context of the extra hydrophobic methionine at its N-terminus (added for translation initiation of our VSV-G chimaera) as compared to the WT protein of the mutant virus. Of note, additional mutations were recorded in the FT-resistant virus strains, especially in the capsid, membrane and envelope proteins (**Fig. 7C**), known to be important in virus attachment and entry, and presumably in viral fitness which can also contribute to overcome drug pressure. Additional studies of those mutants could yield a better understanding of the observed FT resistance.

A more clear resistance pattern was obtained in HCT116 cells that were exposed to high concentrations of FT (**Fig. 6**). From our study, it was clear that the A70V mutation in Sec61α had the strongest impact on FT escape. This mutation is positioned at the plug domain at the lumenal side of Sec61α, and has, to our knowledge, not been reported for other Sec61 inhibitors. However, the adjacent residue S71 has been identified as one of the mutated AA in resistance studies for decatransin [61], coibamide A [62] and mycolactone [63]. The contribution of the A70V mutation to FT resistance was verified in DENV-infected cells, in transfected cells with the DENV-VSV chimaera, but also in a cell-free *in vitro* translation assay. In the latter we could assess FT resistance at the specific level of DENV protein translocation across Sec61, thus, excluding additional effects of other putative (random) mutations in various proteins in the HCT116 cells that might directly or indirectly affect virus replication. The unique A70V mutation suggests that FT might position in or interact with Sec61α in a slightly different way, or bind less strongly to the translocon as compared to CK147 [45]. This would support a high client selectivity of the compound with little impact on other proteins (**Suppl. Fig. 6B**). A more detailed proteomic study will be required to further profile the client spectrum of FT.

To provide a structural model, molecular docking was used to position FT in the Sec61 pore using the Sec61:CK147 complex conformation characterized by cryoEM [45] (see supplement for details; CK147 was removed from the coordinate file before docking FT). The most favorable docking pose for FT (see **Fig. 8**) closely overlapped with that of CK147. This may have resulted in part because the CK147-bound structure was used as docking template. By structural superposition with 8B6L.pdb, we also positioned the DENV2 C14 SP at the lateral gate. In this orientation, the closest distance between FT and DENV2 is 10.6 Å. The conformational stability of this arrangement was monitored during two independent molecular dynamics simulations of 200 ns length each. In both simulations, DENV2 and FT remained close to their modelled starting conformations (see **Fig. 8** and **Suppl. Fig. 9**). This molecular modelling study hence confirms that FT and the DENV2 C14 SP can be well simultaneously accommodated in the Sec61 pore and suggests that the cryoEM binding pose for CK147 may be a suitable pose for FT as well.

**Figure 8.**
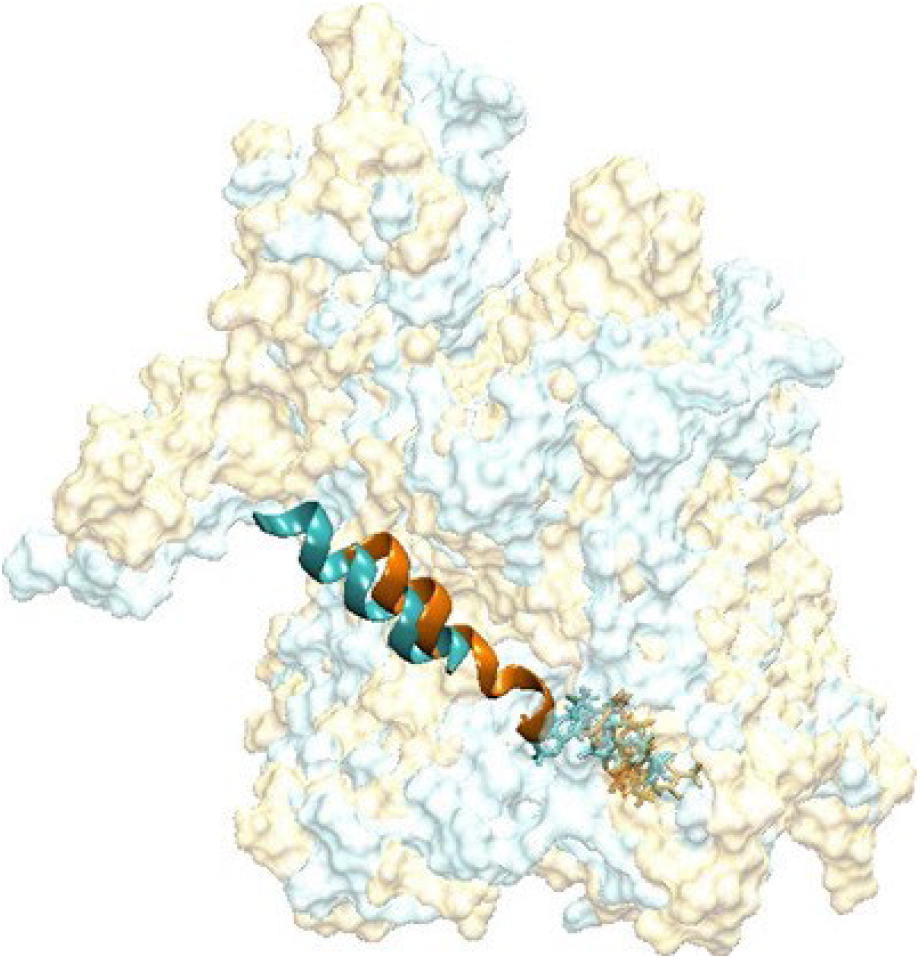
Structural superposition of snapshots from MD simulation #1 (MD1) of SP and inhibitor bound Sec61 at t= 0 ns (cyan) and at t = 200 ns (orange). Sec61 is shown in surface, SP (DENV2 C14) in cartoon and FT is shown in stick representation, respectively.

A big advantage of FT over other described general Sec61 inhibitors (such as mycolactone and Apratoxin A) is its limited toxicity, most likely resulting from a high client selectivity as was seen for CADA [49]. Interestingly, another CADA-analog (VGD040) was recently reported to selectively downmodulate the multi-pass thyrotropin receptor in the context of Graves hyperthyroidism without apparent toxicity in mice. The effectivity and safety of this drug *in vivo* underlines the possibility for clinical applications of CADA analogs designed to target specific SPs [64]. Given the high conservation of Sec61α among species, and the consistent antiviral activity of FT in both host and vector, *in vivo* DENV studies in *Aedes* mosquitos could be justified for a potential antiviral application of FT in vector control settings.

In conclusion, this study shows that a small molecule can selectively interfere with the Sec61-dependent translocation process of flavivirus polyproteins and confirms the critical role of ER translocation in the replicative cycle of flaviviruses. The results from this work on FT open up new therapeutic avenues for antivirals and contribute to our pandemic preparedness efforts.

## EXPERIMENTAL PROCEDURES

### Compounds

Flavitransin (FT; formerly VGD020) was purchased from TCG Lifesciences (India). It was synthesized according to a reported synthesis scheme for VGD020 [51]. Synthesis of FT returned an additional minor compound fraction of an inactive form of FT that was designated as FT_IN_ (See **Supplementary**). Compounds were dissolved in dimethyl sulfoxide (DMSO) and stored as a 10 mM stock at room temperature.

### Cells

African green monkey kidney cells (Vero; American Type Culture Collection; ATCC, Manassas, VA, USA) were grown in Dulbecco’s Modified Eagle’s medium (DMEM; Thermo Fisher Scientific, USA) supplemented with 10% fetal bovine serum (FBS, Hyclone, Cytiva, MA, USA) and 2 mM L-glutamine (Thermo Fisher Scientific). Human hepatocarcinoma cells (Huh7) were cultured in DMEM supplemented with 10% FBS, 1 mM sodium pyruvate (Thermo Fisher Scientific), 0.01 M HEPES (Thermo Fisher Scientific) and 1 X non-essential amino acids (NEAA, Thermo Fisher Scientific). Human embryonic kidney epithelium cells (HEK293T, ATCC) were grown in DMEM supplemented with 10% FBS. Chinese hamster ovaria epithelium cells (CHO-K1, ATCC) were grown in HAM’s F-12 medium (Thermo Fisher Scientific) supplemented with 10% FBS. Baby hamster kidney cells (BHK-21; ATCC) were cultured in Minimum essential medium (MEM, Thermo Fisher Scientific) supplemented with 10% FBS and 2 mM L-glutamine and 0.075% sodium bicarbonate (Thermo Fisher Scientific). Wild-type Raji/0 and Raji/DC-SIGN^+^ were kindly provided by Dr. Geijtenbeeck [65]. The cell lines were cultivated in RPMI-1640 supplemented with 10% FBS and 2 mM L-glutamine. HCT116 cells (ATCC) were cultured in McCoy’s 5A medium (Thermo Fisher Scientific) supplemented with 10% FBS. Finally, C6/36 cells (ATCC) were cultivated in Leibovitz’s L-15 (Thermo Fisher Scientific) supplemented with 10% FBS, 0.01M HEPES and 1X NEAA.

Human peripheral blood mononuclear cells (PBMCs) were prepared from fresh buffy coats (Belgian Red Cross) of healthy donors by standard density gradient centrifugation over Lymphoprep (Nycomed, Oslo, Norway). Monocytes were isolated from the PBMCs and differentiated into immature MDDC by incubation with 20 ng ml^-1^ IL-4 and GM-CSF (Peprotech, London, United Kingdom) for 5 days [66].

All cell lines were maintained at 37°C in a humidified 5% CO_2_ incubator except for larva mosquito cells from *Aedes albopictus* C6/36, which were cultured at 28 °C in the absence of CO_2_. Cell were passaged routinely every three to four days.

### Viruses

Several laboratory and clinical strains of DENV and ZIKV were used in this study. Laboratory-adapted DENV1 Djibouti strain D1/H/IMTSSA/98/606 (GenBank accession number AF298808.1) [67], DENV3 H87 prototype strain (M93130.1) [68], and DENV4 Dak strain HD 34460 (MF004387.1; unpublished data) were kindly provided by X. de Lamballerie (Université de la Méditerranée, Marseille, France). The YFV-17D Stamaril vaccin strain was kindly provided by Sanofi Pasteur (Lyon, France). The laboratory-adapted DENV2 New Guinea-C strain (NGC; VR-1584) and the prototype ZIKV strain MR766 (MK105975.1; VR-84) were purchased at ATCC. DENV1 to 3 clinical isolates were obtained by Kevin Ariën from the Institute of Tropical Medicine (ITM; Antwerp, Belgium). All samples for routine diagnostics from patients presenting at the ITM polyclinic are stored after completion of the routine tests. The ITM has a policy that sample leftovers of patients presenting at the ITM polyclinic can be used for research unless the patients explicitly state their objection. The Institutional Review Board of ITM approved the policy of this presumed consent, as long as patients’ identity is not disclosed to third parties. DENV was isolated from stored serum samples from previously diagnosed acute dengue infections.DENV and ZIKV viral strains were generated by propagation in C6/36 mosquito cells. Titration of the virus was performed by plaque assay using BHK-21 cells [69]. DENV1, DENV3 and DENV4 did not induce viral plaques and therefore the viral titers were determined through flow-cytometry according to the literature [70]. YFV was passaged in Vero cells and titrated on Huh7 cells.

### Antiviral assays

Vero, Huh7 and HCT116 cells (2.0 x 10^5^ cells/ml) were seeded and, after overnight attachment, infected with DENV2. All laboratory-adapted strains were used at MOI 0.5, except for DENV2 and ZIKV in Vero cells and HCT116 cells and YFV in Huh7 which were used at MOI 1. Clinical isolates were used at an MOI of 1. Immature MDDCs (2.5 x 10^6^ cells/ml, 24-well plates, Corning, New York, USA) were infected with DENV2 NGC at MOI 0.5. Unbound virus was removed after 2 h followed by incubation of the infected cells with FT or FT_IN_ .The supernatant was collected 3-5 days post infection and viral RNA was quantified by quantitative reverse transcription-PCR (RT-qPCR). MDDCs were lysed and intracellular viral RNA was quantified by RT-qPCR. In case of YFV, virus-induced cytopathogenic effect (CPE) was assessed by the MTS method.

### Time-of-drug-addition assay

Monolayers of Vero cells (2.5 x 10^5^ cells/ml, 96 -well plates) were infected with DENV2 NGC (MOI of 3) or with ZIKV MR766 (MOI of 1). The virus inoculum was removed after 2 h and replaced by infection medium after washing of the cells with DPBS. FT (1 µM) was added to the infection medium either 30 min prior to infection (-30’), at the time of infection (0 hpi) or at 4, 8, 12, 16, 20 and 23 hpi. The entry inhibitors dextran sulfate with a MW of 40000 Da (DS, 4 µM) or Laby A1 (10 µM) and the nucleoside analogue NITD 008 (10 µM; Tocris Bioscience, Bristol, UK) were included as reference compounds. At 24 hpi, the cells were lysed, and intracellular viral RNA copies were quantified by RT-qPCR.

### RNA isolation and RT-qPCR

For laboratory-adapted strains, intracellular viral RNA or RNA in the supernatant was extracted and quantified with the Cells Direct One-Step RT-qPCR kit (Thermo Fisher Scientific) according to the manufacturer’s instructions. Viral copy numbers were quantified using 5 µl of RNA and 15 µl Master Mix following the standard cycling program, that is reverse transcription at 50°C for 15 min, denaturation at 95°C for 2 min, and then 40 cycles of amplification at 95°C for 15 s and 60°C for 1 min. In the case of clinical isolates and DENV2 from HCT116 infection, viral RNA was first extracted from the supernatant using the QIAamp Viral RNA Mini kit (Qiagen, Hilden, Germany) and quantified using 15 µl of TaqMan Fast Virus 1-Step Master Mix (Applied Biosystems) with specific primer/probe set in addition to 5 µl of viral RNA under the standard cycle conditions: reverse transcription at 50°C for 5 min, denaturation at 95°C for 20 s, and then 40 cycles of amplification at 95°C for 15 s and 60°C for 1 min. RT-QPCR was performed using the QuantStudio 5 Real-Time PCR System (Applied Biosystems, Massachusetts, USA). The primers and probes were used at final concentrations of 900 nM and 250 nM, respectively. The primer and probe sequences were obtained from the nucleotide sequence of the 3’ UTR of DENV1-4 (Genbank sequence AF298808.1, M29095.1, M93130.1, MF004387.1) and the E gene of ZIKV (Genbank sequence MK105975.1) [69] using Snapgene software and are described in Supplementary Table 1. The sequence of the 3’UTR of each viral strain was confirmed by Sanger sequencing (Macrogen Europe, BA Amsterdam, The Netherlands). Viral copy numbers were calculated based on a standard curve of serial 10-fold dilutions of viral DNA template with known copy numbers. All data were analysed with QuantStudio Design & Analysis Software (v1.5.0; Applied Biosystems) and visualised in GraphPad Prism 10.4.1 software.

### Analysis of protein expression in DENV infected cells

Monolayers of huh7 cells (3 x 10^5^ cells/ml, 6-well plates) were treated with 5-fold serial dilutions of FT (0-2 µM) or FT_IN_(10 µM) and after two hours of pre-incubation infected with DENV2 at MOI 0.3. Unbound virus was washed away 2 hpi and cells were further incubated in presence of the compounds. Cells were lysed 48 hpi in NP-40 buffer (50 mM Tris pH 8.0, 150 mM NaCl, 1% NonidetP-40 (Thermo Fisher Scientific), NP-40), supplemented with 0.4 mM/µM phenylmethylsulfonyl fluoride (PMSF, Thermo Fisher Scientific) and Complete protease inhibitor cocktail (Roche, Bazel, Switzerland). The lysates were centrifuged (17.000 g for 10 min at 4°C) and analysed by immunoblotting.

### Plasmids and mutagenesis

The pXJ construct encoding C_14_-prM-E was kindly provided by D. Hoepfner (Novartis Institute for BioMedical Research, Basel, Switzerland). The simian virus 5 (V5, GKPIPNPLLGLDST), 3 FLAG (DYKDHDGDYKDHDIDYKDDDDK) and 2 MYC (EQKLISEEDLNGEQKLISEEDL) epitopes were introduced in the C_14_-prM-E plasmid via mutagenesis with primers elongated with the tags. To create the pXJ construct expressing E_23_-NS1-V5, E_23_-NS1 was isolated from DENV2 NGC strain cDNA via PCR reaction with specific primers (IDT) and cloned into a PCR linearized pXJ vector encoding a V5 epitope. For flow cytometry experiments, the VSV-G protein (from VSV-G expressing pMD2.G plasmid, Didier Trono Lab, Addgene) was inserted into a pCAGGS-tGFP-P2A-BFP expression vector after linearization of the vector via EcoRV cleavage (New England Biolabs, Ipswich, Massachusetts, USA). The signal peptide of VSV-G in the pCAGGS-tGFP-P2A-BFP expression vector was exchanged with the DENV/ZIKV targeting sequences by assembly of their PCR fragments. To generate DENV resistance C14 mutant plasmids, PCR fragments were generated with specific primers surrounding the mutations. The N-GLYC-DV2-C_14_-prM_8_-pPL constructs were generated by cloning of the PCR amplified C_14_-prM_8_ fragment from the pXJ expression vector into the linearized pGEM4 vector encoding N-GLYC-Streptag-(pPL). All PCRs were performed with the Q5 Hot Start HF DNA polymerase (New England Biolabs) according to the manufacturer’s instructions. PCR products were purified from the agarose gel using the Nucleospin Gel and PCR clean-up kit (Macherey Nagel, Düren, Germany). Substitutions were performed using the Q5 site-directed mutagenesis kit (New England Biolabs) according to the manufacturer’s instructions. Inserts were integrated in the expression vectors based on overlapping DNA ends with the NEBuilder HiFi DNA assembly kit (New England Biolabs) following the manufacturer’s instructions. NEB5α-competent *Escherichia coli* cells (New England Biolabs) were transformed with the generated plasmid DNA, as described by the manufacturer’s protocol. Next, the bacteria were inoculated on lysogeny broth (LB) agar plates supplemented with the appropriate selection antibiotics. After ON incubation, single colonies were picked and inoculated in liquid LB medium for ON growth at 37°C. Finally, plasmid DNA was isolated from NEB5 α using the Nucleospin Plasmid Transfection grade system (Machar-Nagel), supplemented with an endotoxin removal wash. The concentration of all constructs was determined with a NanoDrop 1000 spectrophotometer and sequences were confirmed by automated capillary Sanger sequencing (Macrogen Europe).

### Transient transfection

HEK293T, HCT116 (400 000 cells/ml) or CHO-K1 (200 000 cells/ml) cells were seeded one day prior to transfection. The cells were transiently transfected with 2.5 µg plasmid using DNA Lipofectamine LTX & PLUS Reagent (Invitrogen, Thermo Fisher Scientific), according to the manufacturer’s instructions.

Six hours post transfection, FT (0-0.4-2-10 µM) or FT_IN_ (10 µM), together with 200 nM of MG 132 (Sigma, Merck, Burlington, MA, USA) in case of transfection with DV2-C_14_-prM-E, E_23_-NS1-V5 and 2K-NS4B-V5 encoding plasmids, was added. Cells transfected with DV2-C_14_-prM-E (WT or mutant), E_23_-NS1-V5 and 2K-NS4B-V5 encoding plasmids were collected 24 h post-transfection. The collected cells were lysed in NP-40 buffer and DENV protein expression was analysed by immunoblotting as described below. Endoglycosidase H treatment was performed according to the manufacturer’s protocol after cell lysis (Promega, Wisconsin, USA). HEK293T cells transiently transfected with the chimeric tGFP-P2A-BFP plasmid constructs were collected 6 hours post transfection and transferred to 96-well plates filled with 5-fold serial dilutions of FT (0-10 µM) and FT_IN_(0-10 µM). After a compound treatment of 18 hours, these cells were collected and tGFP and BFP expression was measured by flow cytometry as described below.

### Viability assay’s

All cytotoxicity assays were performed on a monolayer of cells (2.0 x 10^5^ cells/ml) by incubation with 2-fold serial dilutions of FT or FT_IN_, starting at higher concentrations than in antiviral assays (range 0 to 80 µM). After 3-5 days, the viability of cells in the presence of the compounds was assessed by adding 0.2 % MTS [3-(4,5-dimethylthiazol-2-yl)-5-(3-carboxymethoxyphenyl)-2-(4-sulfophenyl)-2*H*- tetrazolium; Promega, Leiden, The Netherlands] for 2 h incubation at 37°C. The optical density was measured at 490 nm with the SpectraMax Plus 384 (Molecular Devices, USA) and CC_50_ values (compound concentration reducing the number of cells by 50%) were calculated based on the absorbance of negative (*i*.*e*. cells without compound) and positive (*i*.*e*. culture medium) control samples. The ability of HCT116 cells to proliferate in the presence of FT (0-50 µM) was assessed as described in [45]. In contrast to the standard protocol in which compound treatment is started after overnight attachment of the cells, an alternative protocol was included in which the compounds were administered at the time of seeding.

### Cell-free in vitro translation and translocation

The HotStarTaq DNA polymerase kit (Qiagen) was used to amplify and linearize DNA of interest (full length prM or the first 122 amino acids of the E protein, preceded by their targeting sequences (C_14_, M_21_, respectively)) from the plasmid (from pXJ expression vectors) using 2 steps PCR to include translation initiation sites. PGEM4 expression vectors, which include T7 polymerase initiation site, were linearized by NHE-I (New England Biolabs) restriction digestion. cDNA fragments for the truncated N-GLYC-C_14-_pPL nascent chain were generated by PCR with specifically designed primer sets. PCR fragments were purified with the NucleoSpin Gel and PCR Clean-up Kit (Macherey Nagel) and transcribed *in vitro* using T7 RNA polymerase (RiboMAX system, Promega). RNA was purified using the Nucleospin RNA clean-up kit (Macherey Nagel). mRNA transcripts were translated in rabbit reticulocyte lysate (Promega) or in cell lysate derived from insect (*Spodoptera frugiperda*) cell cultures (RTS 100 Insect Membrane Kit; Westburg) in the presence of L-^35^S-methionine (Perkin Elmer, Waltham, MA, USA). Translations were performed at 30°C in the presence or absence of ovine pancreatic microsomes (RM), FT or FT_IN_, supplemented with RNasin (Promega). In addition, proteinase K (Roche) digestion assays were performed on ice for 30 min and quenched by administration of PMSF (Thermo Fisher Scientific). Samples were washed with low-salt buffer (80 mM KOAc, 2 mM Mg(OAc)_2_, 50 mM HEPES, pH 7.6) and radiolabelled proteins were isolated by centrifugation (10 min at 21382*g*, 4 °C). Radiolabelled proteins were separated by SDS-PAGE on 10% or 4-12% Criterion XT Bis-Tris gels (Bio-Rad, Hercules, CA, USA) in MES buffer (Bio-Rad). The separated proteins were detected by phosphor imaging (Cyclone Plus phosphor storage system, Perkin Elmer) and quantified with the accompanying Image Quant TL software.

### Immunoblotting

NP40 lysates were boiled in reducing sample buffer (120 mM Tris·HCl pH 6.8, 4% SDS, 20% glycerol, 100 mM dithiothreitol, 0.02% bromophenol blue) and analysed by immunoblotting as described in [60]. Primary antibodies used to detect native DENV proteins or DENV proteins modified with the respective epitope tag are summarized in Suppl. Table 2. Differences in loaded protein concentrations between lanes were compensated by normalizing for clathrin heavy chain expression. Unconjugated antibodies were detected by binding with HRP conjugated swine anti-rabbit or goat anti-mouse (secondary antibodies, Suppl. Table 2). Secreted NS1 proteins (including a C-terminal V5-tag) were captured from cell supernatant on V5-Trap Agarose beads according to the manufacturer’s protocol (Chromotek, Planegg, Germany) before immunoblotting. Pull down experiments with lysate of CHO- K1 cells transfected with C_14_-prM-E plasmid were performed on V5-Trap Agarose beads like described in the manufacturer’s protocol.

### Flow cytometry

Vero and Huh7 cells (3.0 x 10^5^ cells/ml) were infected with DENV2 at MOI 0.5. C6/36 cells (6.0 x 10^5^ cells/ml) and Raji-0/Raji-DC-SIGN^+^ cells (1.0 x 10^6^ cells/ml) were infected at MOI 0.2. After washing of unbound virus, cells were treated with FT (0-10 µM) or FT_IN_ (10 µM). Cells were collected by after 48- 72h of incubation in presence of the compounds before staining. Raji cells were first stained extracellular with PE labeled rat anti-CD209 antibody (Suppl. Table 3). All infected cells were permeabilized using the BD Cytofix/Cytoperm Fixation/Permeabilization kit (BD Biosciences, San Jose, CA, USA) as described in the manufacturer’s protocol and stained with APC-linked monoclonal mouse anti-DENV type II antibody (Suppl. Table 3). APC fluorophores were linked to the anti-DENV type II antibody with the APC Conjugation Kit-Lightning Link (Abcam, Cambridge, UK) in advance. Receptor expression was analysed with flow cytometry after extracellular staining of MT-4 cells treated with FT or FT_IN_ for 24h. For stainings, cells were washed with ice-cold PBS containing 2% FBS and incubated with antibody dilutions (Suppl Table 3) for 30 min at 4°C in the dark. Excess antibody was removed by washing the cells with DPBS containing 2% FBS. For the flow cytometric analysis of cellular tGFP and BFP levels, no antibody staining was performed. Instead, cells were collected and washed with DPBS. All cells were finally fixed in DPBS containing 1% paraformaldehyde (VWR Life Science brand, Radnor, PA, USA). All flow cytometry data were collected on BD FACS Celesta Cell Analyzer (BD Bioscience) equipped with BD FACSDiva 8.01 software and analyzed in FlowJo X, version 10. Viable cells and single cells were gated for analysis by exclusion of the cell debris and doublets. To quantify the downmodulating activity of FT, IC_50_ values were calculated in GraphPad Prism 8 software (San Diego, CA, USA) based on four-parameter concentration response curves fitted to the data of replicate experiments. The data are normalized to the untreated control (set at 100%) and the normalized tGFP/BFP ratio was determined. The IC_50_ value represents the compound concentration that resulted in 50% reduction in protein expression.

### Selection and genotyping of FT-resistant DENV

DENV2 NGC was passaged on Huh7 cells (5.0 x10^4^ cells, 24 well plates) under increasing FT pressure, starting at 0.1 µM. In parallel, DENV2 was passaged in the absence of compound or in the presence of the inactive FT_IN_ to check for spontaneous mutations or tissue-culture-adapted mutations. For the weekly passaging of the virus, supernatant was collected and used to infect a fresh monolayer of cells. The remaining supernatant was stored at -80°C. Freshly made compound solutions were added each passage. Compound concentrations were increased as virus escape had developed, tracked between passages via microscopic evaluation of CPE. After 75 passages, resistant DENV was propagated on Huh7 cells to generate DENV stocks. The resistant virus stocks were sequenced and used for antiviral experiments. RNA extraction, cDNA library preparation, and nanopore sequencing were performed as described before [71]. In brief, 20X homemade enzyme buffer (1-M Tris, 100-mM CaCl2, and 30-mM MgCl2, pH 8), 1-μl microccocal nuclease (New England Biolabs) and 2 μl benzonase (Merck-Millipore) were added to each virus sample for viral enrichment. After an incubation of 2 h at 37 °C, enzymatic digestion was stopped by adding EDTA to a final concentration of 10 mM. The samples were then centrifuged at 17.000 g for 3 min, and the resulting supernatant was filtered through a 0.8-µM filter by centrifuging at 17.000 g for 1 min. The resulting filtrate was then subjected to RNA extraction using the Viral RNA mini kit (Qiagen), according to the manufacturer’s instructions. Next, cDNA libraries were generated from the extracted RNA using the SISPA method described by Greninger et al. [72]. Clean- up was performed using 1X AMPure XP beads (Beckman Coulter, Brea, CA, USA) according to the manufacturer’s instructions. Sequencing libraries were prepared using the SQK-NBD114-24 kit (Oxford Nanopore Technologies, UK) according to the manufacturer’s protocol. For each sample, 120-ng cDNA was used as input and ∼100 fmol library was loaded onto the R.10.4.1 flow cell. Sequencing was performed on a GridION and basecalling (and demultiplexing) was performed using Guppy v6 and above. Porechop (github.com/rrwick/Porechop) was used to remove sol-Primer sequences added during cDNA generation. CLC Genomics workbench v22.0.1 was used for mapping of the reads.

### Selection and genotyping of FT desensitized-HCT clones

FT-resistant HCT cells were selected and genotyped following the protocol described by Pauwels et al. with some adaptations [45]. Low densities of HCT116 cells (2.5 x 10^6^ cells) were seeded in T25 flasks in the presence of 10 or 25 µM of FT and incubated at 37°C. The compound medium was replaced every 2 days to support selection of resistant cells. After two weeks, cells that survived the compound pressure were collected and thoroughly diluted as single cells over multiple 6 well plates or large petri dishes. The surviving cells were further incubated at 37 °C and microscopically monitored until single cell colonies were detected. These single cell colonies represented the FT-resistant cells and were isolated by clonal ring selection as described elsewhere [45, 73]. RNA was isolated from 5 x 10^6^ cells using the RNeasy Mini Kit (Qiagen) according to the manufacturer’s protocol. Next, 350 ng of the isolated RNA was reverse transcribed into complementary DNA (cDNA) using random primers from the High-Capacity cDNA Reverse Transcription Kit (Thermo Fisher Scientific). Specifically designed primers were used to amplify Sec61α by PCR with Q5 polymerase (Promega) as described in the manufacturer’s protocol. PCR fragments were purified from agarose gel with the Nucleospin Gel and PCR Clean-up Kit and the amplicon length was checked on 1% agarose gels. Nanopore sequencing and subsequent analysis were performed as described previously [45].

### Preparation of semi-permeabilized cells

Confluent FT-resistant or WT HCT116 cells were harvested from a T75 flask by trypsinisation. The collected cells were kept on ice and resuspended in KHM buffer (110 mM Kac, 2mM MgAc_2_, 20 mM HEPES-KOH, pH=7.2 at 4°C) with trypsin-inhibitor (Merck). The buffer was removed and cells were collected by centrifugation at 2000 rpm, for 3 min at 4°C between different treatment steps. Cells were permeabilized by resuspension in KHM buffer supplemented with digitonin (44 µM/1*10^6^ cells, MP Biomedicals, California, USA) for 5min. Next, cells were further diluted in KHM buffer to prevent additional permeabilization. The cytosol was washed out by resuspension of the semi-permeabilized cells in HEPES-buffer (50 mM Kac and 90 mM HEPES-KOH, pH = 7.2 at 4°C) and incubation on ice for 10 min. To digest endogenous RNA, the cells were treated with nuclease (Thermo Fisher Scientific) and 0.1 µM CaCl_2_ by resuspension in KHM buffer. After 12 minutes of incubation at RT, nuclease was inhibited by addition of 0.4 µM EGTA (Sigma, Merck) and removed by resuspension of the cells in KHM buffer. *In vitro* translations were performed with 120.000 semi-permeabilized cells per sample in presence of rabbit reticulocyte lysate.

### Statistical analysis

Statistical analyses were performed using Graph Pad Prism version 10.4.1 for Windows (Graph Pad Software; www.graphpad.com). Unpaired t-test with Welch’s correction was used for the analysis of the *in vitro* translocation, immunoblot and flow cytometric data. Observed differences were regarded as significant if the calculated P-values were *P ≤.05; **P ≤ .01; ***P ≤.001, ***P ≤ 0.0001.

## Acknowledgments

We thank Piet Maes for sequencing support and Eva Pauwels for sharing protocols and giving advice in translocation experiments.

