## Supplemental Methods and Figures for "Sec61 translocon inhibitor Flavitransin blocks selectively dengue virus polyprotein insertion into the ER membrane with pan-flavivirus antiviral potency"

#### SUPPLEMENTARY

##### NMR analysis $FT_{IN}$

*General.* $^{1}$H NMR spectrum was recorded on Avance 400 (Bruker) spectrometer. Chemical shifts (*δ*) are reported in parts per million (ppm) with $CHCl_3$ resonance peak set at 7.26 ppm. Multiplicities are indicated as s (singlet), d (doublet), t (triplet), m (multiplet). Coupling constants, *J*, are reported in Hertz. $^{1}$H NMR assignments were corroborated by 2D experiment ($^{1}$H-$^{1}$H COSY). High resolution mass analysis (ESI) was performed on LTQ ORBITRAP XL Thermo apparatus.

$^{1}$H NMR (400 MHz, $CDCl_3$): δ 7.72 (m, 4H, ArH), 7.69 – 7.61 (d, 4H, , *J* = 7.9 Hz, ArH), 7.29 (d, *J* = 8.1 Hz, 4H, ArH), 6.97 (m, 4H, ArH), 5.23 (s, 4H, $=CH_2$), 3.86 (s, 6H, $OCH_3$), 3.73 (bs, 8H, $=CCH_2N$), 3.14-2.99 (m, 8H, $SO_2NCH_2$), 2.42 (s, 6H, $ArCH_3$), 2.21 (t, *J* = 7.0 Hz, 8H, $2N(CH_2)_2$), 1.96 (d, *J* = 7.0 Hz, 4H, $NCH_2CH$), 1.72-1.58 (m, 12H), 1.57-1.47 (bs, 8H, $NCH_2CH_2CH_2N$), 1,30-1.21 (m, 2H, $NCH_2CH$), 1.22-1.12 (m, 4H), 0.80-0.69 (m, 4H). HRMS (ES+) *m/z* calcd for $C_{62}H_{91}N_6O_{10}S_4^+$ 1207.56795 $[M+H]^+$ found 1207.56605 (5%); $[M+2H]^{2+}$ 604.28789 found 604.28752 (100%).

##### Molecular Docking and Molecular Dynamics

The PDB structure of Sec61 was retrieved from protein data bank entry 8B6C[1] (https://www.rcsb.org/structure/8b6c). Preparation of the receptor structure included adding polar hydrogens and assigning partial atomic charges. The ligand (FT) PDBQT file was generated using AutoDock tools. Docking of FT in the Sec61 pore was conducted using the AutoDock Vina v1.2.3 software package[2,3] with AutoDock Tools v1.5.6[4]. Pymol software was used for visualization. As docking area, a grid box was used of size 20×20×45 $Å^3$ along the Sec61 pore. The exhaustiveness of global search was maintained at 16 and a grid spacing of 0.375 Å was used. All other docking parameters were set at default values. First, the validity of the docking protocol was tested by removing CK147 from 8B6C and redocking it into the Sec61 pore. After superimposition, this gave a RMSD of 0.020 Å for CK147 and a binding score of -12.82 kcal/mol reflecting the suitability of the docking protocol for this class of compounds.

When docking FT in the same manner, the pose with highest binding affinity closely overlapped with that of CK147 in the cryoEM structure (binding score -8.83 kcal/mol). Hence, this pose with highest binding affinity was considered for subsequent MD simulations. The signal peptide of Dengue Virus (DENV2: MTAGMIIMLIPTVMA) was modeled into an alpha-helical conformation using UCSF Chimera v1.17.3.[5] PDB entry 8B6L[6] from Forster's lab was used to position the SP (DENV2) at the lateral gate. The closest distance between modelled SP and docked FT was 10.64 Å. For atomistic MD simulations of the system, the modelled Sec61:SP:FT complex was embedded in a POPC bilayer membrane using CHARMM-GUI web portal.[7]

CHARMM36 force field was employed for proteins[8] and lipids,[9] and TIP3P[10] model for water. The system was solvated with water molecules and neutralized with counterions. The solvated system was energy minimized using the steepest descent method for 5000 steps. The equilibration of the solvated system was conducted in six-steps (the first two in the NVT ensemble, the last four in the NPT ensemble) based on the standard CHARMM-GUI equilibration protocol. The Nosé-Hoover[11] thermostat with a coupling constant of 1.0 ps at 303.15 K and Parrinello-Rahman[12] barostat at 1 bar were used to maintain temperature and pressure of the system. Short-range van der Waals interactions were calculated using a cutoff of 1.2 nm, and long-range electrostatic interactions were treated with the particle-mesh Ewald (PME) method.[13] The LINCS algorithm was used to constrain bond lengths to their equilibrium values. Subsequently, two independent MD simulations (200 ns each) were conducted using the GROMACS (version 2023.2)[14] package, starting with different random velocities to characterize the conformational dynamics of the protein-ligand complex and the potential impact of ligand binding on the Sec61 structure and stability. During the MD simulations, ligand and SP remained stably bound. Structural superpositions of initial (at t= 0 ns: cyan) and final (at t = 200 ns: orange) snapshots are shown in Figures 1 and 2. Here, Sec61 is shown in surface, SP in cartoon and FT in stick representation, respectively.

[4] Morris, G. M.; Huey, R.; Lindstrom, W.; Sanner, M. F.; Belew, R. K.; Goodsell, D. S.; Olson, A. J. AutoDock4 and AutoDockTools4: Automated Docking with

Selective Receptor Flexibility. J. Comput. Chem. 2009, 30 (16), 2785– 2791, DOI: 10.1002/jcc.21256.

[5] E. F. Pettersen, T. D. Goddard, C. C. Huang, G. S. Couch, D. M. Greenblatt, E. C. Meng, T. E. Ferrin, UCSF Chimera—A visualization system for exploratory research and analysis. J. Comput. Chem. 25, 1605–1612 (2004).

[6] Gemmer M, Chaillet ML, van Loenhout J, Cuevas Arenas R, Vismpas D, Gro¨llersMulderij M, Koh FA, Albanese P, Scheltema RA, Howes SC, Kotecha A. Visualization of translation and protein biogenesis at the ER membrane. Nature. 2023 Feb

2;614(7946):160-7.

[7] Jo S, Kim T, Im W. Automated builder and database of protein/membrane complexes for molecular dynamics simulations. PloS one. 2007;2(9):e880. pmid:17849009

[8] Best RB, Zhu X, Shim J, Lopes PE, Mittal J, Feig M, et al. Optimization of the additive CHARMM all-atom protein force field targeting improved sampling of the backbone $\varphi$, $\psi$ and side-chain $\chi1$ and $\chi2$ dihedral angles. Journal of chemical theory and computation. 2012;8(9):3257–3273. pmid:23341755

[9] Klauda JB, Venable RM, Freites JA, O'Connor JW, Tobias DJ, Mondragon-Ramirez C, et al. Update of the CHARMM all-atom additive force field for lipids: validation on six lipid types. The journal of physical chemistry B. 2010;114(23):7830–7843.

pmid:20496934

[10] Jorgensen WL, Chandrasekhar J, Madura JD, Impey RW, Klein ML. Comparison of simple potential functions for simulating liquid water. The Journal of chemical physics.

1983;79(2):926–935.

[11] Evans DJ, Holian BL. The nose–hoover thermostat. The Journal of chemical physics. 1985;83(8):4069–4074.

[12] Parrinello M, Rahman A. Polymorphic transitions in single crystals: A new molecular dynamics method. Journal of Applied physics. 1981;52(12):7182–7190.

[13] Darden T, York D, Pedersen L. Particle mesh Ewald: An N · log (N) method for Ewald sums in large systems. The Journal of chemical physics. 1993;98(12):10089-10092.

[14] Abraham, M. J.; Murtola, T.; Schulz, R.; P´all, S.; Smith, J. C.; Hess, B.; Lindahl, E. GROMACS: High Performance Molecular Simulations through MultiLevel Parallelism from Laptops to Supercomputers. SoftwareX 2015, 1–2, 19– 25, DOI: 10.1016/j.softx.2015.06.001

### SUPPLEMENTARY FIGURES

#### Supplementary Fig. 1

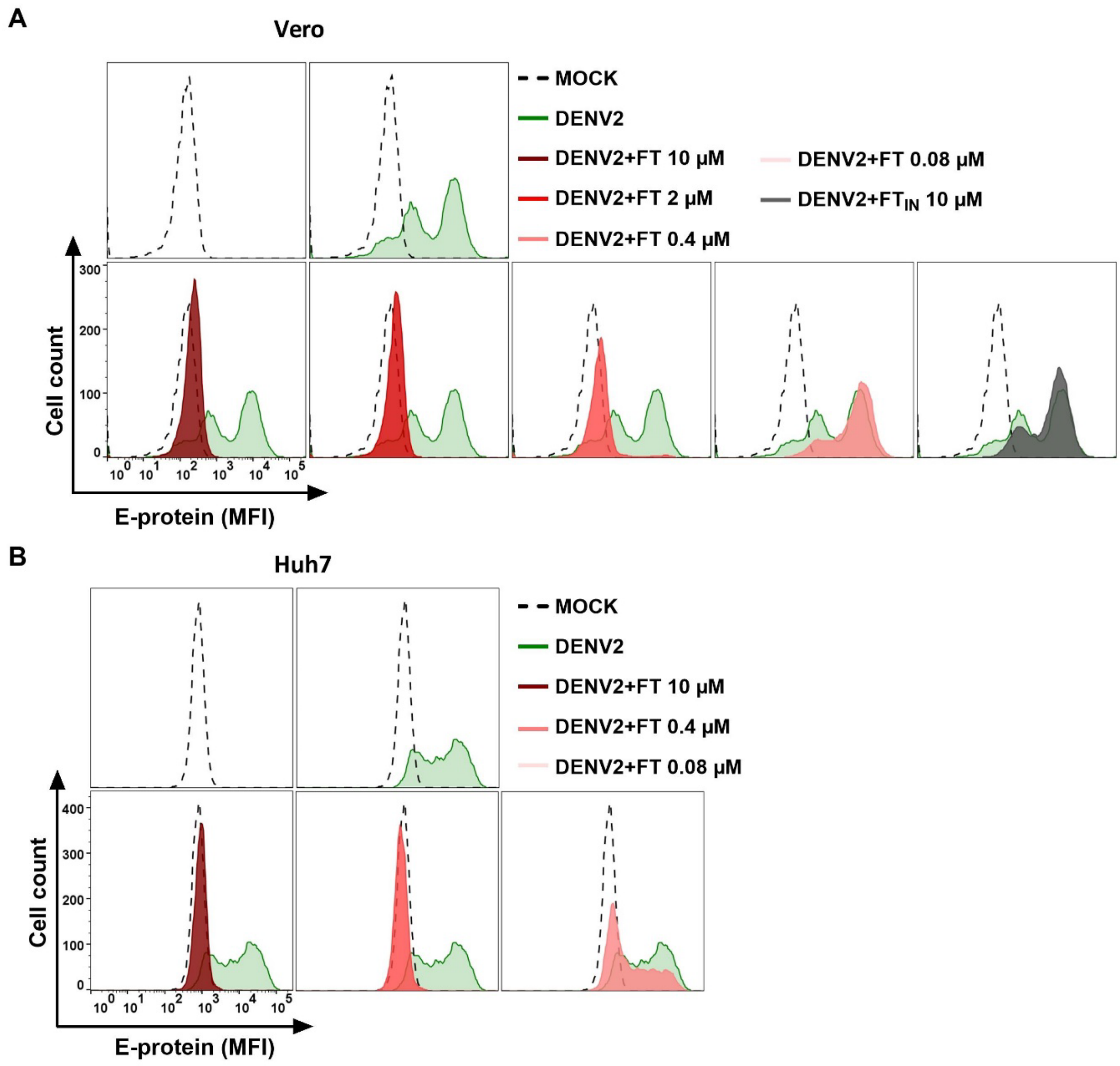


**Suppl. Fig. 1 Potent anti-DENV2 activity of FT in Vero and Huh7 cells.** Viral E protein detection in DENV2 NGC (MOI 0.5) infected (A) Vero and (B) Huh-7 cells, treated with FT or 10 µM $FT_{IN}$ for 72h. Flow cytometric histogram plots show mean fluorescence intensity (MFI) values for E protein, acquired from at least 5,000 cells. Mock-infected cells are in black dotted line, virus infected control cells are in green line, FT treated infected cells in red and $FT_{IN}$ infected cells in grey. Representative histogram plots are given for one out of two experiments. See Fig. 1 for the FT 2 µM and $FT_{IN}$ conditions in Huh7 cells.

#### Supplementary Fig. 2

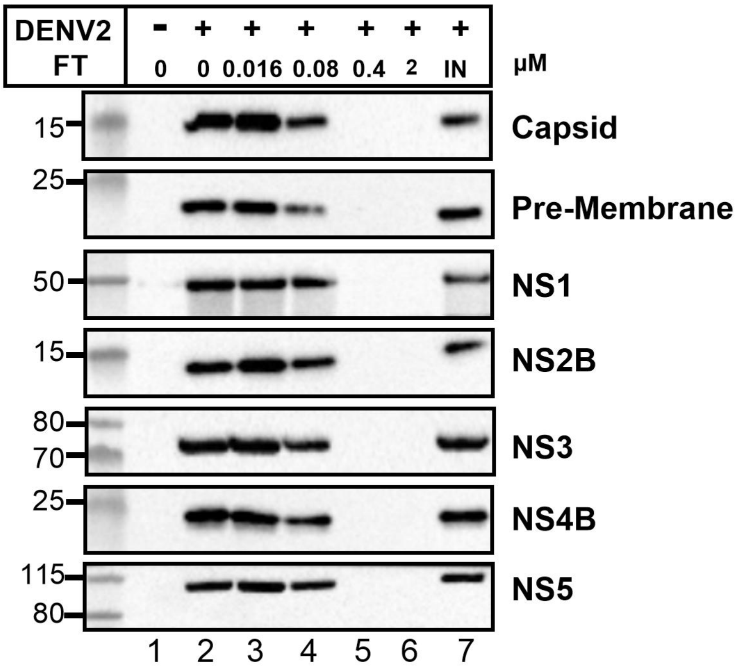


**Suppl. Fig. 2 FT inhibits the expression of DENV2 structural and non-structural proteins in infected Huh7 cells**. Same as in Fig. 1F. Viral protein detection in DENV2 NGC (MOI 0.3) infected Huh7 cells, treated with FT or 10 µM $FT_{IN}$ (indicated as 'IN') for 48h. Same cell lysate samples of Fig. 1F were used for immunoblotting with antibodies against the separate DENV proteins. See Fig. 1F for immunoblots of E and clathrin (loading control).

### Supplementary Fig. 3

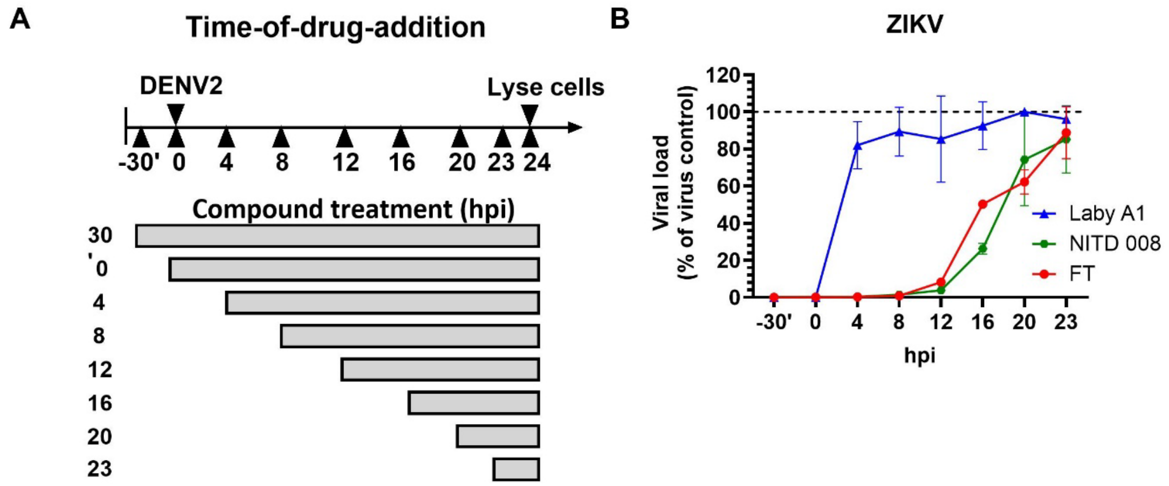


**Suppl. Fig. 3 FT interferes with a post-entry event of the ZIKV life cycle. A,** Schematic representation of the time-of-drug-addition assay. **B,** Time-of-drug-addition assay as explained in Fig. 1H but for infection of Vero cells with ZIKV and Laby A1 (10 µM) as entry inhibitor. Values are mean ± SD; n=3.

#### Supplementary Fig. 4

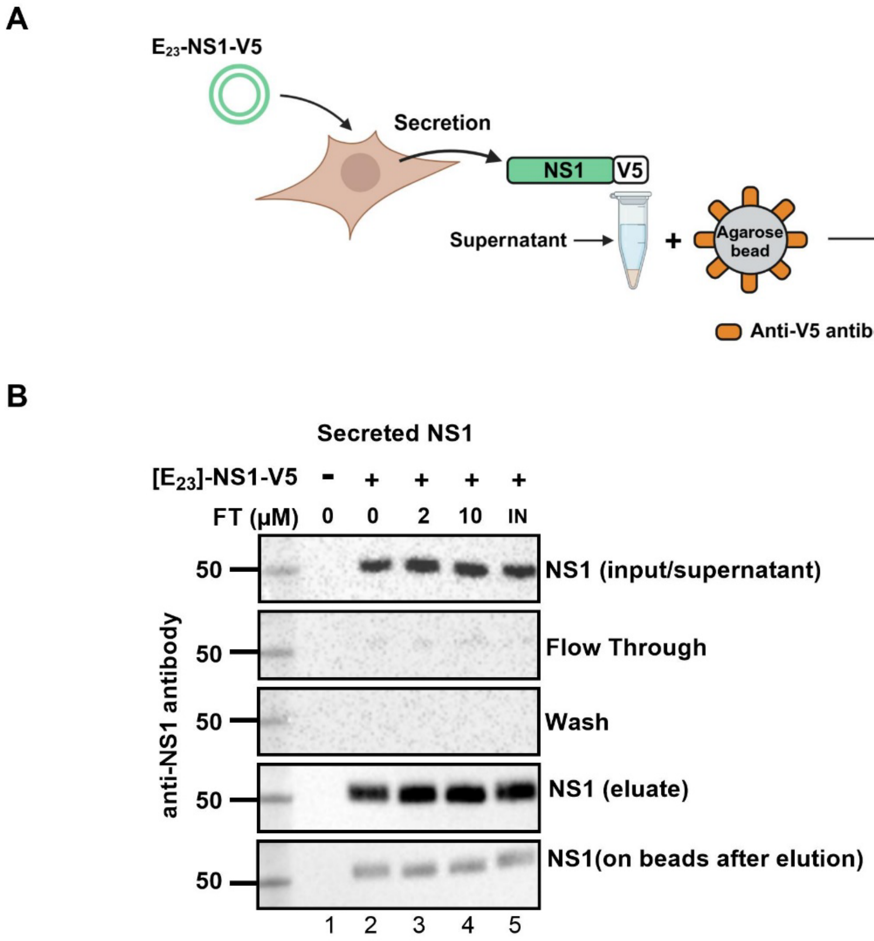


**Suppl. Fig. 4 FT has no effect on the secretion of DENV2 NS1 protein. A,** The supernatant from cells transfected with the $E_{23}$-NS1-V5 plasmid from Fig. 2C was incubated with magnetic agarose beads coated with V5 nanobodies to capture/pull-down V5-tagged secreted NS1. Cartoon created with BioRender (2025); www.BioRender.com. **B,** Extracellular secreted NS1 in supernatant of transiently transfected HEK293T cells (lane 2-5). Immunoblots by detection with an NS1 antibody of (i) input (NS1 in supernatant before pull down), (ii) flow through unbound NS1 after incubation with beads, (iii) wash of beads, (iv) eluted NS1, and (v) NS1 remaining on beads after elution by boiling in sample buffer. One representative blot out of two experiments is shown.

#### Supplementary Fig. 5

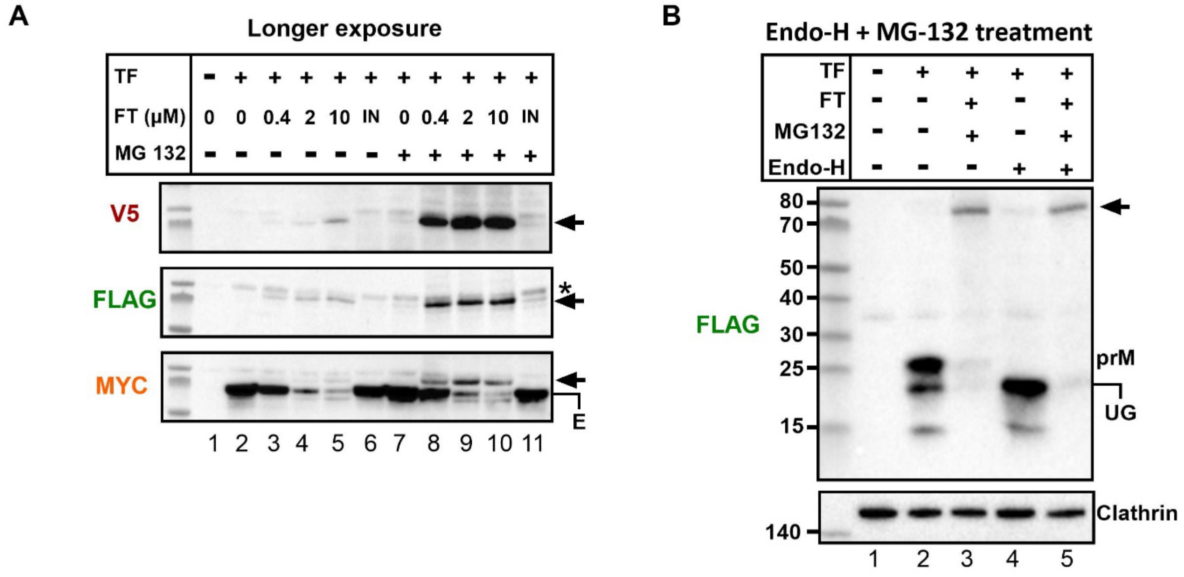


**Suppl. Fig. 5 FT diverts the untranslocated $C_{14}$-prM-E precursor to the proteasomal degradation pathway. A,** Same blots as in Fig. 4B but for longer chemiluminescence exposure times (>saturation) to visualize the precursor (arrow). **B,** Cell lysate of transfected cells (TF) from Fig. 4B treated with FT and MG132 was incubated with Endo-H and subjected to immunoblotting with an anti-FLAG antibody and anti-clathrin (loading control). UG: unglycosylated. The uncleaved precursor (arrow) is not glycosylated.

#### Supplementary Fig. 6

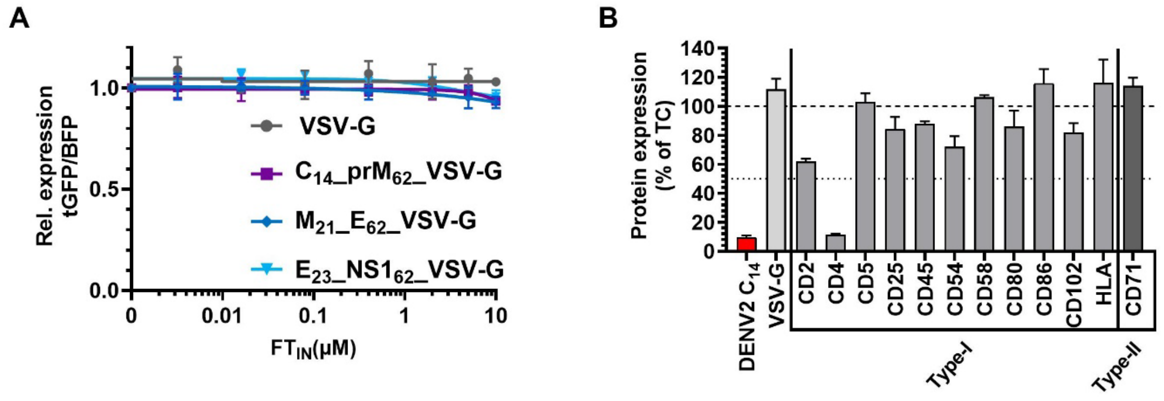


**Suppl. Fig. 6 Flavitransin has a high selectively for the membrane-spanning domain of the DENV capsid subunit. A,** Cells transiently transfected with constructs from Fig. 5A but treated with different concentrations of $FT_{IN}$. GFP and BFP expression (mean fluorescent intensity) was quantified by flow cytometry and normalized to untreated transfection control. Graph shows the four-parameter concentration-response curves of $FT_{IN}$ for the tGFP:BFP ratio. Values are mean ± SD; n=3. **B,** MT-4 cells were treated with FT for 24 hours and the endogenous expression of several cell surface receptors was quantified by flow cytometry using fluorescently labelled (PE/FITC-) antibodies. Bar graphs show the relative expression (mean fluorescent intensities) at 2 µM FT, normalized to the endogenous expression levels in untreated controls (TC). Bars are mean ± SD; 3≤n≤5.

### Supplementary Fig. 7

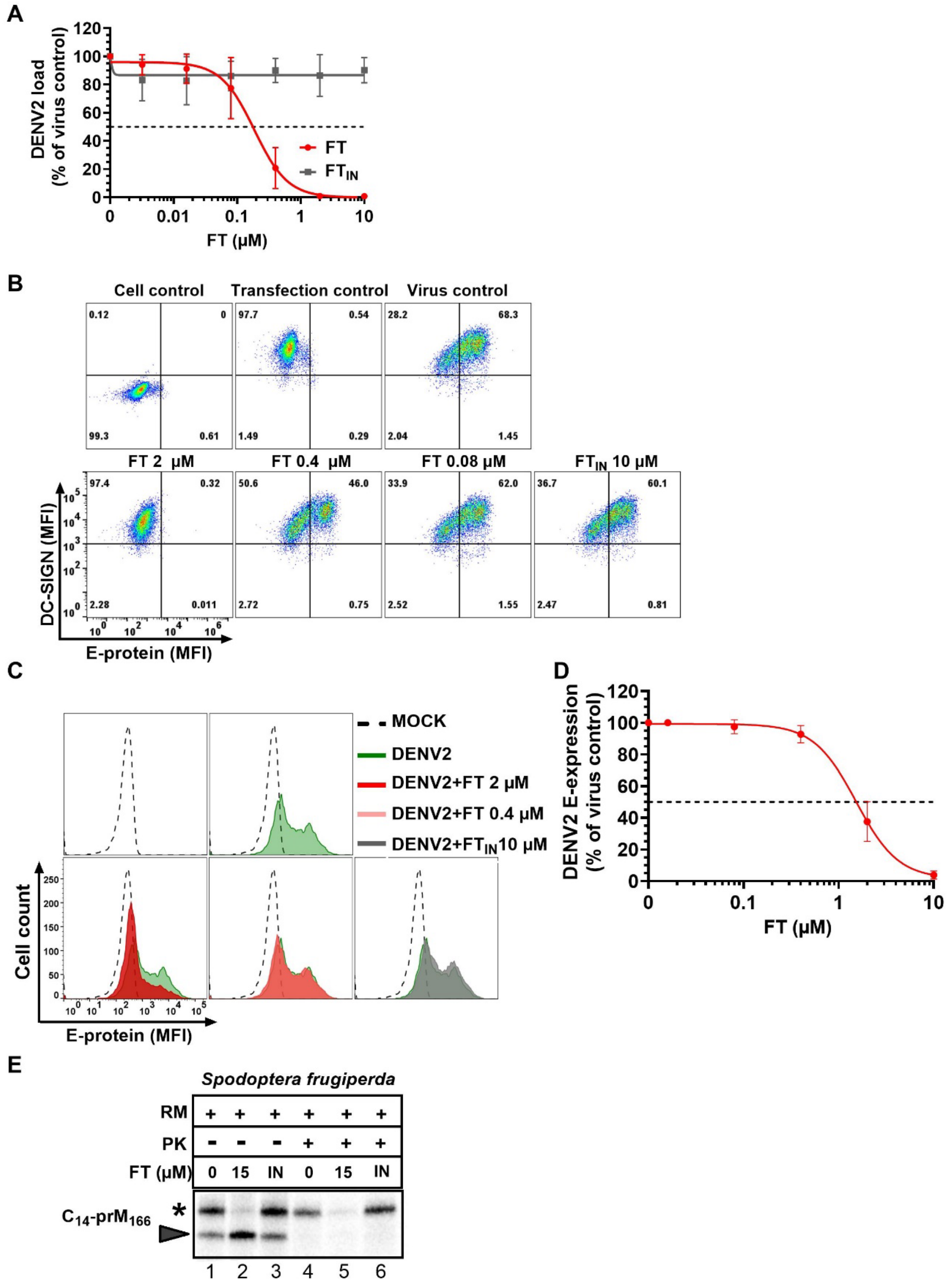


**Suppl. Fig. 7 FT exerts consistent anti-DENV2 activity in different cell types. A,** Four parameter concentration-response curve of FT or $FT_{IN}$ for virus yield in DENV2 NGC-infected (MOI 0.5) monocyte-derived dendritic cells (MDDCs). MDDCs were lysed 48 hpi and viral RNA copy number was quantified by RT-qPCR. Graphs represent the viral load as normalized to the viral copy number of the virus control. Values are mean ± SD of 3 independent experiments with MDDCs derived from 5 different donors. **B,** Raji-DC-SIGN$^+$ cells (stably transfected with the DENV entry receptor CD209) were infected with DENV2 (MOI 0.2) and incubated with FT or $FT_{IN}$ for 48h. Cells were fixed, stained extracellular with anti-CD209 and stained with an anti-DENV2 E antibody after permeabilization. Cell control: infected WT-Raji (Raji-0) cells, Transfection control: uninfected Raji-DC-SIGN$^+$ cells, virus control: infected Raji-DC-SIGN$^+$ cells, FT: infected Raji-DC-SIGN$^+$ cells treated with FT, and $FT_{IN}$: infected Raji-DC-SIGN$^+$ cells treated with 10 µM $FT_{IN}$. Representative dot blots are given for one out of three experiments. Dot blots show MFI values for E and DC-SIGN protein, acquired from at least 8,000 cells per experiment. **C,** C6/36 cells (derived from larva of *S. albopictus*, the principal vector of DENV transmission) were infected with DENV2 (MOI 0.2) and incubated with FT or $FT_{IN}$ for 72h. Flow cytometric histogram plots show MFI values for E protein, acquired from at least 5,000 cells. Mock-infected cells are in black dotted line, virus infected control cells are in green line, FT treated infected cells in red and $FT_{IN}$ infected cells in grey. Representative histogram plots are given for one out of three experiments. **D,** Four parameter concentration-response curve show the relative E-protein expression from (C) as normalized to virus control; values are mean ± SD, n=3 **E,** Transcripts encoding DENV2 $C_{14}$-$prM_{166}$ were translated in a cell free *in vitro* assay using cell lysate of *Spodoptera frugiperda* in the presence of 15 µM FT or $FT_{IN}$ and [$^{35}$S]-labeled methionine, and treated with PK (similar to Fig.3B). Samples were separated by SDS-PAGE and analyzed by autoradiography. Solid arrowhead: unprocessed nascent protein; asterisk: glycosylated (translocated) mature protein. Of note, as this translation mixture also contains the microsomes (RM), translation only of the preprotein cannot be assessed.

#### Supplementary Fig. 8

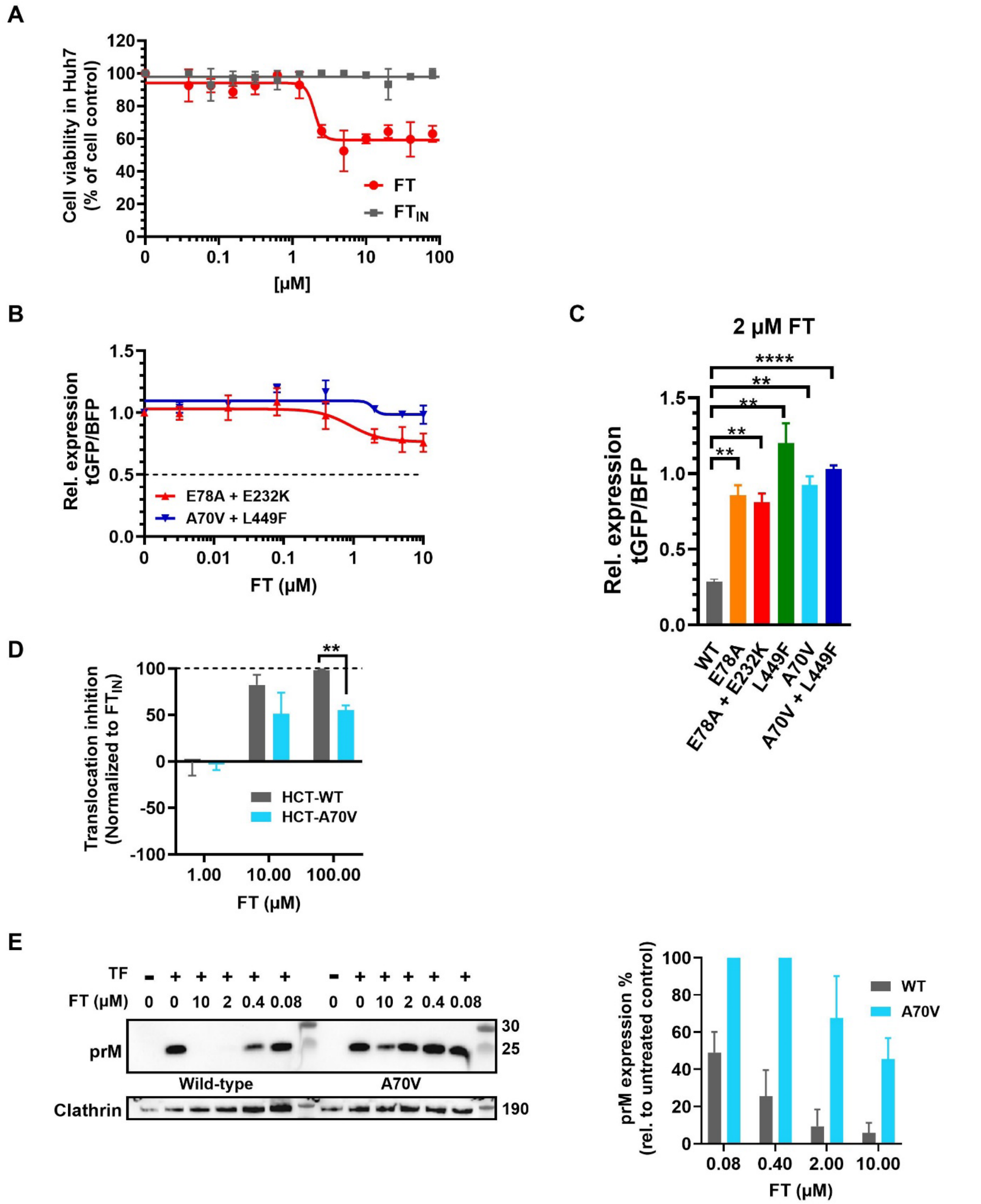


**Suppl. Fig. 8 FT-resistant cells contain mutations in Sec61α and show reduced sensitivity to FT in suppressing DENV prM expression. A,** Four parameter concentration-response curve of FT or $FT_{IN}$ for viability of Huh7 cells. Cytotoxic effects were determined at day 5 post treatment by MTS/PMS assay. Data are normalized to untreated cell controls. Graph shows mean values ± SD, n=2. **B,** Same as in Fig. 6D but for the transfection of the DENV2 $C_{14}$-

$prM_{62}$-VSV-G plasmid constructs with dual Sec61α mutations. **C,** Bars represent the relative tGFP:BFP expression ratio for the 2 µM treatment of FT shown in panel B and in Fig. 6D from three independent experiments. Values are mean ± SD; n = 3. **P<0.01, ****P<0.001, compared to the expression in WT HCT116 cells by multiple unpaired t-test with Welch's correction. **D,** Graphs show the relative translocation inhibition calculated from the translocated protein fraction (asterisk) in fig. 6E normalized to and subtracted from the translocation of the $FT_{IN}$ control. Bars are mean ± SD; n = 3. Multiple unpaired t-tests with Welch's correction were performed to define statistical significant differences with the HCT116 WT control (**p<0.01). The 10 µM samples did not differ significantly. **E,** WT and FT-resistant HCT116 cells were transiently transfected with the $C_{14}$-prM-E construct (from Fig. 4A) and treated with FT for 18 h. Cells were lysed in NP-40 buffer and analyzed by immunoblotting with an anti-V5 antibody. For the cell loading control, an anti-clathrin antibody was used. A representative immunoblot is shown on the left. Bar graph on the right represents relative prM expression levels from clathrin-corrected samples, normalized to untreated transfected controls from three independent experiments. Bars are mean ± SEM, n=3.

#### Supplementary Fig. 9

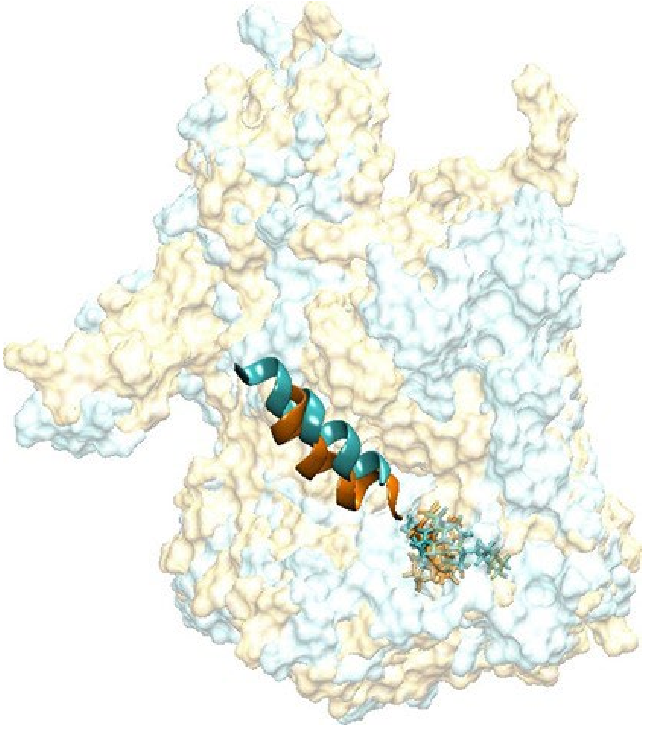

**Suppl. Fig. 9** Structural superposition of initial and final snapshots from MD simulation #2 (MD2) of SP and inhibitor bound Sec61 at t= 0 ns (cyan) and at t = 200 ns (orange). Sec61 is shown in surface, SP (DENV2 C14) in cartoon and FT is shown in stick representation, respectively.
